# iPSC culture expansion selects against putatively actionable mutations in the mitochondrial genome

**DOI:** 10.1101/2020.11.05.369694

**Authors:** Maike Kosanke, Colin Davenport, Monika Szepes, Lutz Wiehlmann, Tim Kohrn, Marie Dorda, Jonas Gruber, Kaja Menge, Maike Sievert, Anna Melchert, Ina Gruh, Ulrich Martin

**Affiliations:** Leibniz Research Laboratories for Biotechnology and Artificial Organs (LEBAO), Department of Cardiothoracic-, Transplantation and Vascular Surgery, REBIRTH - Research Center for Translational Regenerative Medicine, Hannover Medical School, 30625 Hannover, Germany; Biomedical Research in Endstage and Obstructive Lung Disease (BREATH), Member of the German Center for Lung Research (DZL), 30625 Hannover, Germany; Research Core Unit Genomics, Hannover Medical School, 30625 Hannover, Germany

## Abstract

While human induced pluripotent stem cells (iPSCs) offer fascinating prospects for research and clinics, evaluating their genomic stability before applications is of utmost importance. During reprogramming, clonal hiPSC lines derived from the same parental cell population were observed to harbor different mitochondrial DNA (mtDNA) variants and variant heteroplasmy levels. It is unknown to date to which extent this unequal segregation of heteroplasmies between cells arises from mosaicism in the parental cell population, selection on mutated mtDNA molecules on the cellular or organelle level, genetic drift during reduction of mtDNA during reprogramming (genetic bottleneck), or *de novo* mutations. We analyzed mtDNA variants in 26 clonal iPSC lines by mtDNA sequencing. We did not observe a strong bottleneck or any selection for cells with specific mtDNA variants (clonal eliteness) during reprogramming. iPSC culture expansion, however, selects against putatively actionable mutations in the mitochondrial genome that may affect mitochondrial function and cell metabolism. In contrast, heteroplasmy levels of neutral variants remained stable or increased within clonal iPSC lines during culture expansion. Interestingly, the mtDNA copy number per cell got transiently reduced during targeted differentiation of iPSCs into cardiomyocytes, but mtDNA heteroplasmy levels were not affected. Altogether, our results point towards a scenario in which intra-cellular selection on mtDNA during culture expansion, but not during reprogramming or differentiation, pivotally shapes the mutational landscape of the mitochondrial genome in iPSCs. Other mechanisms such as inter-cellular selection, genetic bottleneck, and genetic drift exert minor impact.

## Introduction

The availability of human induced pluripotent stem cells (iPSCs) ^1^ offers fascinating prospects for disease modelling and the development of tailored cellular therapies. However, on the path toward clinical application one major hurdle is the concern for genomic instability of iPSCs that may cause loss of function and tumorigenic potential of their derivatives ^2–4^. Therefore, the evaluation of effects and possible biological significance of genetic changes for research use and patient safety is of utmost importance. The majority of research is focusing on the investigation of genomic stability of the nuclear genome. However, a part of a cell’s genetic information, including 13 proteins of the electron transport chain (ETC) essential for oxidative phosphorylation (OXPHOS) and 24 RNAs, is encoded in the mitochondrial genome ^5^. Mutations of those highly conserved genes are the cause of a variety of human diseases, especially affecting tissues with high energy demand such as skeletal muscle, the myocardium or the nervous system, and age-related conditions ^6,7^.

The mitochondrial genome exhibits a polyploid set with a few hundred up to thousands of 16.6kb large circular mitochondrial DNA (mtDNA) molecules that are continuously replicated independently of the cell cycle (relaxed replication) ^5^. The existence of multiple mtDNA copies per cells allows a phenomenon called heteroplasmy which describes the simultaneous existence of wild-type and mutated mtDNA molecules in a cell. Conversely, the state when only one mtDNA genotype is present in a cell is defined as homoplasmy. However, the mtDNA composition and mitochondria network are not fixed but subjected to permanent flux. Therefore, the heteroplasmy level of mtDNA variants within a cell can change over time, but during cell division the segregation of mutated and wild-type mtDNA molecules to cell progenies can also be unequal. Hence, it is not surprising that during reprogramming clonal iPSC lines derived from the same parental cell population were observed to harbor different variants and heteroplasmy levels ^8–14^. This unequal segregation of heteroplasmies between cells during reprogramming might arise from three sources acting on an inter-cellular and intra-cellular level with different forces depending on the external conditions, namely i) *de novo* mutations ^8,10,15^, ii) genetic mosaicism in parental cell population leading to genetically distinct iPSC clones, and iii) random allele drift during genetic bottleneck ^13,14,16^. Several recent studies report nuclear reprogramming as cause of *de novo* mtDNA mutations in iPSCs ^8,10,15^. However, Payne et al. ^17^ introduced the concept of ‘Universal Heteroplasmy’, demonstrating that mosaicism of heteroplasmic mtDNA variants in somatic cells, albeit at low levels (< 1%), appears to be a universal finding among healthy individuals ^17^. Recently, Deuse et al. could confirm via single-cell sequencing of somatic cells that the heteroplasmy levels of variants obtained in bulk sequencing masks a wide range of heteroplasmies in the individual cells ^15^. Accordingly, many reports show evidence that the majority if not all mtDNA variants in iPSCs are pre-existing in individual somatic parental cells and arise from this mosaicism in the corresponding parental cell population ^13,14^. Some mtDNA variants were pre-existing apparently at a very low frequency within the batch of parental cells but up to 100-fold enriched in a clonal iPSC line during reprogramming ^13,15^.

During reprogramming, somatic mitochondria are largely replaced by immature mitochondria resembling organelle morphology and distribution in embryonic stem cells (ESCs), and mitochondrial metabolism is switching towards glycolysis ^18–20^. During this process, mtDNA copy number per cell is reduced which is assumed to lead to a genetic bottleneck ^21,22^. Previous studies modeling the effect of bottlenecks showed that such a reduction in mtDNA pool increases the effect of random genetic drift on mtDNA segregation within a cell and between cell offspring during cell division ^23,24^. However, at the same time this bottleneck might expose mutations to selective forces, and subtle selective pressures can exert maximal impact ^16,21,23,25^. As of yet, it is not understood to what extent the segregation is driven by random allele drift or selection ^14^. In contrast to the nuclear genome, selection on mutated mtDNA molecules can act on an inter-cellular or intra-cellular level. On the cellular level, genetically encoded inequalities in cell fitness of parental cells can lead during reprogramming to eliteness of cells to attain iPSC state and their dominance in the reprogramming niche ^26,27^. However, high mutational burden or pathogenic mutations in specific mtDNA regions, such as non-coding transcript regions for tRNAs and replication or translation start sites which lead to loss of physiological integrity and higher reactive oxygen species (ROS) production, hinder reprogramming ^9,10,16,28–30^. On intra-cellular level, certain mtDNA variants can be selected ^29^. As mtDNA molecules are uniparentally inherited and the mutation rate is 6-20 fold higher than that of genomic DNA ^31,32^, theoretically, without a repair mechanism, mtDNA would continuously accumulate mutations ultimately ending in a “mutational meltdown” called Muller’s ratchet ^16,33^. To avert this, during female germline development and early embryogenesis fragmentation of mitochondria and mitochondrial selective autophagy (mitophagy) purges mutated mtDNA and mitochondria ^16,29,33–36^. As iPSC experience rejuvenation to an ESC-like state ^19^, a similar mechanism might act during reprogramming which leads to mitophagy of damaged organelles or reduced segregation of such ^16^.

After reprogramming, genetic drift and selection continue to form the mutational landscape of the mitochondrial genome during prolonged culture of iPSCs ^11–15,37^. Although most studies report that mtDNA variants did not experience any substantial change of heteroplasmy level during long-term culture, there is also evidence for both positive ^14,15^ and negative selection of mtDNA variants ^11–13^, without clear comprehension of the causes or mechanisms. On one hand, mitochondrial heteroplasmic variants were observed to arise in or dominate human iPSC and ESC lines during prolonged culture ^14,15,37^. Notably, single cell analysis has revealed heterogeneous heteroplasmies among cell populations and there were no indication of selection against potentially pathogenic variants during culture ^14^. On the other hand, various studies reported that extended passaging of iPSC clones with high level of heteroplasmy affecting specific genes led to reduction of mutant mtDNA ^11–13,21,29^.

It is, however, currently unknown to what extent the diverse and contradictory mechanisms, such as clonal eliteness, random genetic drift, positive selection and clearance or repair of mutated mtDNA, shape the mutational landscape of mtDNA. We have therefore evaluated to what extent and at which stage random genetic drift, selection, and protective mechanism for averting accumulation of mtDNA variants might have an impact in iPSCs. To this end, we have analyzed 26 clonal iPSC lines and their corresponding parental cell populations by mtDNA sequencing and determination of their copy number to gain insight into the dynamics and mechanism behind mtDNA segregation during reprogramming, prolonged culture, and differentiation.

## Results and Discussion

### No evidence for clonal eliteness or mutation clearance mechanisms during reprogramming

To analyze the effect of reprogramming on mtDNA segregation and mutational landscape, we generated 26 clonal iPSC lines of early passages of endothelial cells (EC) of 6 neonatal (human umbilical vein EC; hUVEC) and 4 aged donors (human saphenous vein EC; hSVEC; age 64-88 years). To assess inter-clonal differences, 3 iPSC clones were derived per donor, with exception of D#6 hUVEC and D#40 hSVEC, for which only one clone each was analyzed. In contrast to prior assumptions, mtDNA copy number in early passage iPSCs (p4-9) was, with, on average, 1252 mtDNA copies per cell, similar to ECs of the corresponding parental cell populations (~1114 copies per cell at ~ p4) and ESCs (~934 mtDNA copies per cell) (Fig. 1a). Although there was no distinct reduction and, hence, bottleneck in mtDNA copy number in iPSCs compared to their parental cells, the mitochondria network was remodeled to a more immature state with mitochondria being punctual and located in the perinuclear region (Fig. S1) as previously described ^19^.

**Fig. 1:**
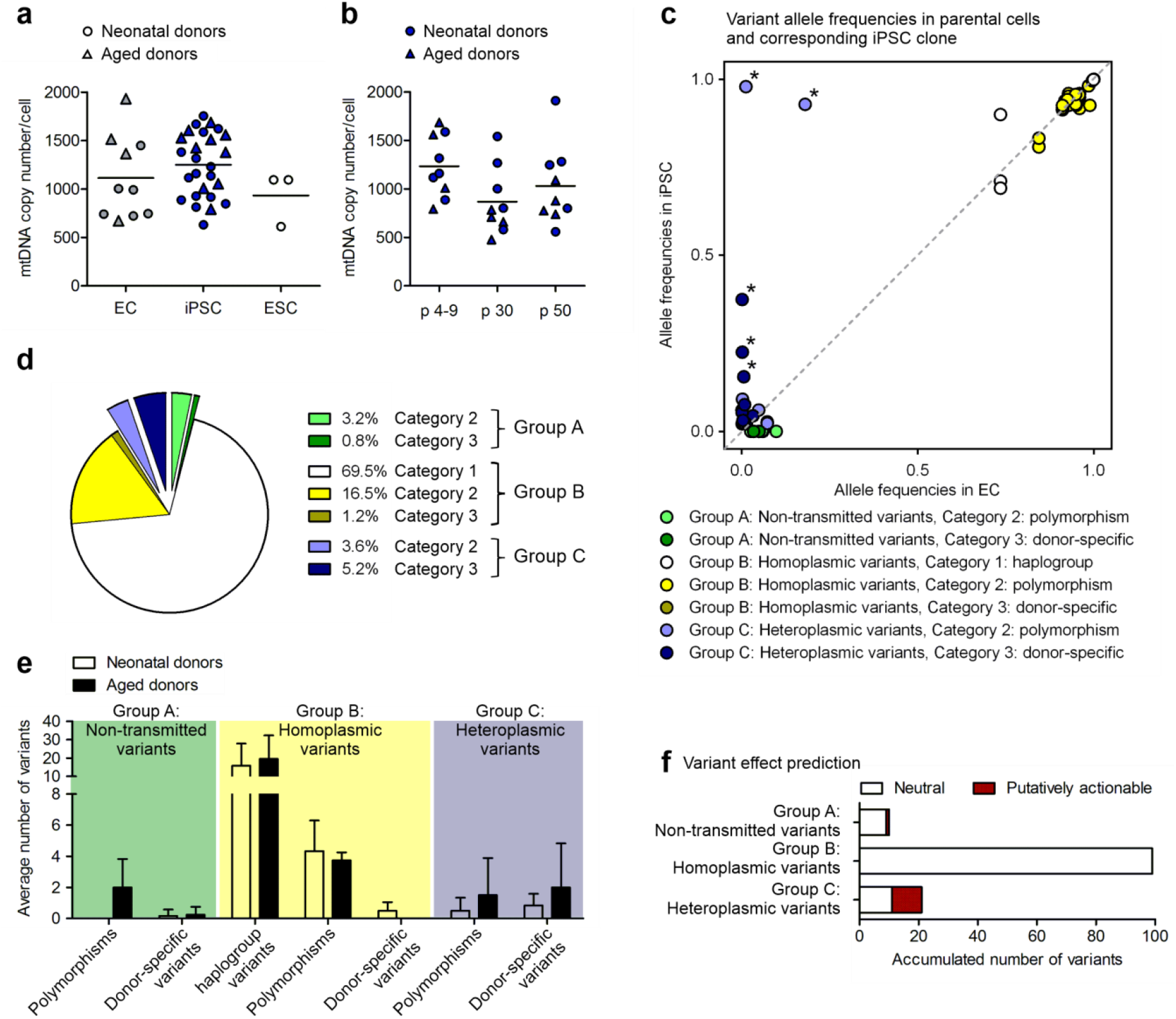
Characterization of mtDNA copy numbers and mtDNA variants in iPSCs and their parental cells. **a** Average mtDNA copy number per cell in parental endothelial cells (EC), EC-derived iPSC clones and ESC lines. EC N = 10 donors; iPSC N = 26 early passage iPSC clones; ESC N = 3 lines; for each N mtDNA copy number was measured as 1-5 biological replicates. **b** Average mtDNA copy number per iPSC clone in different passages of culture expansion. N = 4-5 iPSC clones; 1 biological replicate. **c** Corresponding allele frequencies (AF) of all variants in iPSC clones and parental cell populations detected by mtDNA sequencing of 26 early passage iPSC clones and their corresponding 10 parental cell populations. Group A: non-transmitted variants were detected only in parental cell populations (AF > 0.02). Group B: homoplasmic variants in parental cell population and iPSCs derived thereof. Group C: heteroplasmic variants detected in an iPSC clone with AF > 0.02 but less frequent in parental cell population. Category 1 comprises haplotype variants, category 2 polymorphisms defined by a population frequency (MITOMAP (NCBI GenBank)) ≥ 0.02 and category 3 variants that are specific to a donor, but rare in the global human population. **d** Proportion of variant groups and categories of all variants. **e** Average number of variants per group and category in iPSC clones and/or corresponding parental cell population of neonatal and aged donors. Neonatal donors N = 6 donors with 16 iPSC clones; Aged donors N = 4 donors with 10 iPSC clones. Displayed is mean with SD. **f** Effect of variant on gene functionality was predicted based on a consensus of in silico prediction algorithms obtained from snpEff impact, CADD, Condel, and HmtVar. Graph displays ratio of variants with neutral and putatively actionable prediction within Group A (non-transmitted), Group B (homoplasmic), and Group C (heteroplasmic).

mtDNA of early passages of iPSCs (mean p6.5) and the corresponding parental cell populations, at a similar passage as subjected to reprogramming (mean p4.5), was sequenced. Analysis of mtDNA variants found in iPSCs and ECs revealed 3 groups. Group A, non-transmitted variants, includes variants detected only within the parental cell populations that were apparently not transmitted to any iPSC clone of the representative donor (detection limit allele frequency (AF) 0.02). Those variants were detected in the parental cell population with AF < 1 and therefore exclusively represent variants present in subpopulations of parental cells (Fig. 1c, Table 1, Table S1). In contrast, groups B and C contain transmitted variants. Group B comprises homoplasmic variants that were detected in iPSC clones and their corresponding parental cell populations. Despite one variant detected in two iPSC clones, group C variants were detected as heteroplasmic mutations in one iPSC clone per donor, only (Fig. 1c, Table 1, Table S1). The heteroplasmies are defined for every variant separately as the percentage of variant allele count per total allele count at the variant locus (= AF*100). Although, in most cases the AF of those variants within the parental cell population was below the general detection limit of 0.02, careful assessment of variants within their genomic context taking local error rates in account allowed us to confirm pre-existence of more than half of the variants with statistical confidence (p-value 0.05) (Table S2) (see Methods). Although for the other half of variants we could not prove pre-existence with statistical confidence (Table S2, indicated by value in brackets), previous reports support our assumption that most of these variants arose from the parental cell population as well ^13,14^.

**Table 1:** mtDNA variants in iPSC clones and corresponding parental cell populations. Number of variants per group and category of all variants detected by mtDNA sequencing of 26 early passage iPSC clones and corresponding 10 parental cell populations (detection limit: allele frequency (AF) > 0.02). Variants of group A, non-transmitted variants, were detected only in parental cell populations. Group B variants were homoplasmic in parental cell population and iPSCs derived thereof. Group C, heteroplasmic variants, were detected in an iPSC clone with AF > 0.02 but were less frequent in the corresponding parental cell population. Haplotype variants are those that define a haplogroup (Category 1). Polymorphism are defined by a population frequency (MITOMAP (NCBI GenBank)) ≥ 0.02 (Category 2) and donor-specific variants were unique to a donor and rare in the population context (Category 3).

| Donor | Haplo-group | Non-transmitted<br>(Group A) |  | Transmitted<br>(Groups B+C) |  |  |  |  |
| --- | --- | --- | --- | --- | --- | --- | --- | --- |
|  |  |  |  | Homoplasmic<br>(Group B) |  |  | Heteroplasmic<br>(Group C) |  |
|  |  | Poly-morphisms<br>(Cat. 2) | Donor-specific<br>(Cat. 3) | Haplogroup<br>variants<br>(Cat. 1) | Poly-morphisms<br>(Cat. 2) | Donor-specific<br>(Cat. 3) | Poly-morphisms<br>(Cat. 2) | Donor-specific<br>(Cat. 3) |
| D#1 hUVEC | H7a1 | 0 | 0 | 9 | 3 | 0 | 0 | 1 |
| D#2 hUVEC | H1c3 | 0 | 0 | 10 | 5 | 1 | 0 | 0 |
| D#3 hUVEC | H2a1c | 0 | 1 | 7 | 3 | 0 | 2 | 2 |
| D#4 hUVEC | J1c8a | 0 | 0 | 28 | 3 | 1 | 1 | 1 |
| D#5 hUVEC | J1b1a3 | 0 | 0 | 34 | 8 | 1 | 0 | 0 |
| D#6 hUVEC | H87 | 0 | 0 | 7 | 4 | 0 | 0 | 1 |
| Neonatal donors | Mean | 0.0 | 0.2 | 15.8 | 4.3 | 0.5 | 0.5 | 0.8 |
| D#31 hSVEC | H1c2 | 4 | 0 | 9 | 4 | 0 | 0 | 2 |
| D#37 hSVEC | H10b | 0 | 0 | 8 | 3 | 0 | 4 | 6 |
| D#38 hSVEC | I3 | 3 | 0 | 32 | 4 | 0 | 1 | 0 |
| D#40 hSVEC | J1c5a | 1 | 1 | 29 | 4 | 0 | 0 | 0 |
| Aged donors | Mean | 2.0 | 0.3 | 19.5 | 3.8 | 0.0 | 1.3 | 2.0 |
| Total | Mean | **0.8** | **0.2** | **17.3** | **4.1** | **0.3** | **0.8** | **1.3** |
|  | Sum | 8 | 2 | 73 | 23 | 3 | 8 | 13 |

Variants of each group can be further categorized: Category 1, haplotype variants, are homoplasmic variants that define the haplogroup of every donor (Table 1, Table S1). Category 2 variants comprise polymorphisms defined by a population frequency of the variant (NCBI GenBank frequency (GB) obtained from MITOMAP database) > 0.02. Category 3 variants are such that are unique to a donor and rare in human populations. Those category 3 variants are, therefore, called hereafter donor-specific variants (Table 1, Table S1).

Group B variants (homoplasmic in iPSC clones and corresponding parental cell population) represent the vast majority of all variants (> 87%) detected in mtDNA. Variants of this group B are mainly haplotype variants or polymorphisms (category 1 and 2) (Fig. 1d, Table 1). Only around 4% (with > 3% polymorphisms (category 2)) of all variants were detected only within the parental cell populations (group A) and almost 9% of all variants in iPSCs were heteroplasmic (group C) variants, which were typically rare among the parental cell population (Fig. 1d, Table 1). The fraction of polymorphisms (category 2) of around 3.5% is similar in group A and C, while, interestingly, donor-specific (category 3) variants account for the majority of variants within group C (Fig. 1d, Table 1). Hence, the number of heteroplasmic donor-specific variants (group C, category 3) is substantially higher than the detected number of non-transmitted donor-specific variants (group A, category 3) (Table 1) suggesting a higher mosaicism within parental cell population below the resolution capacity of bulk mtDNA sequencing.

Remarkably, in contrast to Kang et al. ^10^, but in accordance with Payne et al. ^17^ we did not observe a dependency of overall mitochondrial mutational load of iPSCs with donor age (Fig. 1e, Table 1). However, there was a trend that aged donors contain, on average, a slightly higher number of group A (non-transmitted) and C (heteroplasmic) variants, indicating higher mosaicism in cell source of aged donors (Fig 1e, Table 1) as reported by previous studies ^32^.

Next, we analyzed the functional consequences of variants to evaluate to which extent clonal eliteness or negative selection might impact their transmission from parental cells to iPSC clones. Variant effect prediction based on a consensus of *in silico* prediction algorithms revealed that all group B (homoplasmatic) variants and 90% of group A (non-transmitted) variants are neutral (Fig. 1f). Strikingly, almost 48% of the group C (heteroplasmic) variants in iPSCs are putatively actionable (Fig. 1f), suggesting that mtDNA segregation during reprogramming does not act selectively against potentially damaging mutations. However, while we cannot exclude positive selection, we also found no direct evidence for enrichment of individual mutations or any mutational hot spot in specific genes during reprogramming.

Notably, the heteroplasmy level of the group C (heteroplasmic) variants was generally low (below 10%; AF < 0.1) in iPSCs and the bulk of their corresponding parental cells, with the exception of 5 variants (Fig. 1c, marked with *) that showed increased heteroplasmy levels in single iPSC clones. While 2 of these 5 variants were neutral polymorphisms with very high heteroplasmy levels > 90% (AF > 0.9) in individual iPSC clones, the other 3 variants were putatively actionable donor-specific with intermediate heteroplasmy levels of 16-38% (AF 0.16-0.38) in single iPSC clones, affecting NADH dehydrogenase (ND) 4 and ND5 genes of the ETC Complex I. Interestingly, Complex I, in particular the ND5 gene, is frequently mutated in diverse diseases and cancer ^38,39^. However, our data do not allow any conclusion whether the affected iPSC clones simply represent stochastic events, where iPSC clones were derived from rare parental cell clones with increased heteroplasmy levels, or whether positive selection processes occurred. Anyway, we could not observe altered mitochondrial features or metabolic functionality of the ND5 mutations in affected iPSC clones (D#3 hUVEC C2 with 38% m.13099G>A and D#37 hSVEC C10 with 23% m.12686T>C) (Fig. S2, marked with *) compared to their sister clones (D#3 hUVEC C1 and D#37 hSVEC C4) derived from the same parental cell population. All iPSC clones displayed a similar proliferation rate measured as population doubling, mtDNA copy number per cell, specific yield coefficient of lactate per glucose molecule, mitochondrial content, mitochondrial membrane potential, and ROS content (Fig. S2). As iPSC metabolism is not predominantly based on OXPHOS and as the heteroplasmy rates were 23% and 38%, only, this result is not unexpected and in accordance to observations of previous work ^9,13,21,30^, but also decreases the likelihood of a clonal eliteness on the inter-cellular level during reprogramming.

### Prolonged culture expansion leads to clearance of putatively actionable mtDNA mutations

There are contradictory reports regarding the propagation of mtDNA mutations during prolonged culture. While some studies report increase and dominance ^14,15,37^, other studies observed a reduced proportion of mutated mtDNA during culture ^11–13^. 7 of the iPSC clones that had been analyzed in an early passage, were also sequenced in later passages p30 and p50 to trace the dynamic of mtDNA heteroplasmies during culture expansion.

It is noteworthy that 86 % of the group C variants detected in later passages were also detectable in early iPSC passages and / or the corresponding parental cell population with statistical confidence (p-value 0.05). The remaining variants were apparently present in the parental cells and early passages, too, but could not be confirmed with statistical confidence (p-value 0.05). Hence, similar to reprogramming, there was no evidence for *de novo* mutagenesis during prolonged culture expansion (Table 2).

**Table 2:**
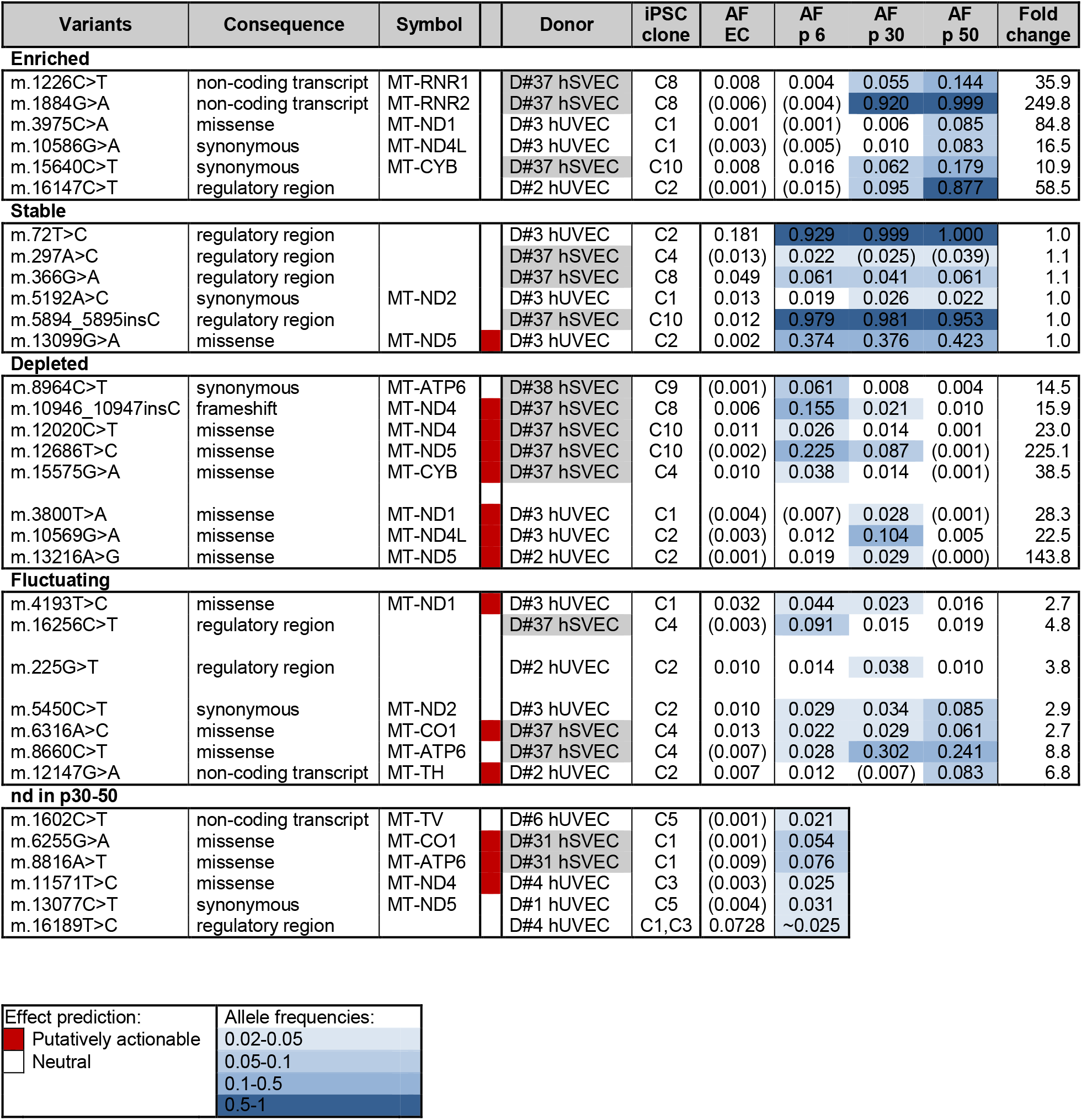
mtDNA variants heteroplasmy levels change during culture expansion. Heteroplasmy levels of iPSC-enriched (Group C) variants were monitored during iPSC culture by mtDNA sequencing of 7 iPSC clones at early (on average p6.5), intermediate (p30), and late passage (p50). Allele frequencies (AF) of variants are displayed for parental cell populations and the different iPSC passages. Shades of blue indicate heteroplasmy levels. Heteroplasmy levels increased during iPSC culture expansion (group of enriched heteroplasmic variants; fold change of variant heteroplasmy > 10), stayed stable during culture expansion (stable; fold change ~1), or decreased (depleted; fold change > 10). Heteroplasmic variants for which fold change of heteroplasmy level did not allow confident classification into one of the above mentioned groups are defined as fluctuating heteroplasmic variants. A last group “not determined (nd) in p30-50” comprises those heteroplasmic variants that were found in the iPSC clones but were only analyzed in early passage (p6). Effect prediction, based on a consensus of *in silico* prediction algorithms obtained from snpEff impact, CADD, Condel, and HmtVar, suggest putatively actionable variants (marked red) and neutral (unmarked) variants. Aged donors are shaded gray. (); existence of variant in parental cell population or iPSC clone not confirmed with statistical confidence (p-value > 0.05).

| Variants | Consequence | Symbol | Donor | iPSC clone | AF EC | AF p 6 | AF p 30 | AF p 50 | Fold change |
| --- | --- | --- | --- | --- | --- | --- | --- | --- | --- |
| **Enriched** |  |  |  |  |  |  |  |  |  |
| m.1226C>T | non-coding transcript | MT-RNR1 | D#37 hSVEC | C8 | 0.008 | 0.004 | 0.055 | 0.144 | 35.9 |
| m.1884G>A | non-coding transcript | MT-RNR2 | D#37 hSVEC | C8 | (0.006) | (0.004) | 0.920 | 0.999 | 249.8 |
| m.3975C>A | missense | MT-ND1 | D#3 hUVEC | C1 | 0.001 | (0.001) | 0.006 | 0.085 | 84.8 |
| m.10586G>A | synonymous | MT-ND4L | D#3 hUVEC | C1 | (0.003) | (0.005) | 0.010 | 0.083 | 16.5 |
| m.15640C>T | synonymous | MT-CYB | D#37 hSVEC | C10 | 0.008 | 0.016 | 0.062 | 0.179 | 10.9 |
| m.16147C>T | regulatory region |  | D#2 hUVEC | C2 | (0.001) | (0.015) | 0.095 | 0.877 | 58.5 |
| **Stable** |  |  |  |  |  |  |  |  |  |
| m.72T>C | regulatory region |  | D#3 hUVEC | C2 | 0.181 | 0.929 | 0.999 | 1.000 | 1.0 |
| m.297A>C | regulatory region |  | D#37 hSVEC | C4 | (0.013) | 0.022 | (0.025) | (0.039) | 1.1 |
| m.366G>A | regulatory region |  | D#37 hSVEC | C8 | 0.049 | 0.061 | 0.041 | 0.061 | 1.1 |
| m.5192A>C | synonymous | MT-ND2 | D#3 hUVEC | C1 | 0.013 | 0.019 | 0.026 | 0.022 | 1.0 |
| m.5894_5895insC | regulatory region |  | D#37 hSVEC | C10 | 0.012 | 0.979 | 0.981 | 0.953 | 1.0 |
| m.13099G>A | missense | MT-ND5 | D#3 hUVEC | C2 | 0.002 | 0.374 | 0.376 | 0.423 | 1.0 |
| **Depleted** |  |  |  |  |  |  |  |  |  |
| m.8964C>T | synonymous | MT-ATP6 | D#38 hSVEC | C9 | (0.001) | 0.061 | 0.008 | 0.004 | 14.5 |
| m.10946_10947insC | frameshift | MT-ND4 | D#37 hSVEC | C8 | 0.006 | 0.155 | 0.021 | 0.010 | 15.9 |
| m.12020C>T | missense | MT-ND4 | D#37 hSVEC | C10 | 0.011 | 0.026 | 0.014 | 0.001 | 23.0 |
| m.12686T>C | missense | MT-ND5 | D#37 hSVEC | C10 | (0.002) | 0.225 | 0.087 | (0.001) | 225.1 |
| m.15575G>A | missense | MT-CYB | D#37 hSVEC | C4 | 0.010 | 0.038 | 0.014 | (0.001) | 38.5 |
| m.3800T>A | missense | MT-ND1 | D#3 hUVEC | C1 | (0.004) | (0.007) | 0.028 | (0.001) | 28.3 |
| m.10569G>A | missense | MT-ND4L | D#3 hUVEC | C2 | (0.003) | 0.012 | 0.104 | 0.005 | 22.5 |
| m.13216A>G | missense | MT-ND5 | D#2 hUVEC | C2 | (0.001) | 0.019 | 0.029 | (0.000) | 143.8 |
| **Fluctuating** |  |  |  |  |  |  |  |  |  |
| m.4193T>C | missense | MT-ND1 | D#3 hUVEC | C1 | 0.032 | 0.044 | 0.023 | 0.016 | 2.7 |
| m.16256C>T | regulatory region |  | D#37 hSVEC | C4 | (0.003) | 0.091 | 0.015 | 0.019 | 4.8 |
| m.225G>T | regulatory region |  | D#2 hUVEC | C2 | 0.010 | 0.014 | 0.038 | 0.010 | 3.8 |
| m.5450C>T | synonymous | MT-ND2 | D#3 hUVEC | C2 | 0.010 | 0.029 | 0.034 | 0.085 | 2.9 |
| m.6316A>C | missense | MT-CO1 | D#37 hSVEC | C4 | 0.013 | 0.022 | 0.029 | 0.061 | 2.7 |
| m.8660C>T | missense | MT-ATP6 | D#37 hSVEC | C4 | (0.007) | 0.028 | 0.302 | 0.241 | 8.8 |
| m.12147G>A | non-coding transcript | MT-TH | D#2 hUVEC | C2 | 0.007 | 0.012 | (0.007) | 0.083 | 6.8 |
| **nd in p30-50** |  |  |  |  |  |  |  |  |  |
| m.1602C>T | non-coding transcript | MT-TV | D#6 hUVEC | C5 | (0.001) | 0.021 |  |  |  |
| m.6255G>A | missense | MT-CO1 | D#31 hSVEC | C1 | (0.001) | 0.054 |  |  |  |
| m.8816A>T | missense | MT-ATP6 | D#31 hSVEC | C1 | (0.009) | 0.076 |  |  |  |
| m.11571T>C | missense | MT-ND4 | D#4 hUVEC | C3 | (0.003) | 0.025 |  |  |  |
| m.13077C>T | synonymous | MT-ND5 | D#1 hUVEC | C5 | (0.004) | 0.031 |  |  |  |
| m.16189T>C | regulatory region |  | D#4 hUVEC | C1,C3 | 0.0728 | ~0.025 |  |  |  |

|  |  |
| --- | --- |
| Effect prediction: | Allele frequencies: |
| Putatively actionable | 0.02-0.05 |
| Neutral | 0.05-0.1 |
|  | 0.1-0.5 |
|  | 0.5-1 |

Investigation of the regions affected by mutations showed that around 60% of all heteroplasmic group C variants were located in coding regions (Fig. S3a, black). Hence, the group C variants were slightly more frequent in coding regions compared to group A (non-transmitted) and group B (homoplasmic) variants (Fig. S3a). Furthermore, compared to group A and B variants which constitute almost entirely nucleotide transitions (transition/transversion ratio (Ts/Tv) > 30), group C variants are characterized by lower T>C transitions compared to C>T and by higher proportion of transversion events (Ts/Tv 3.4) (Fig S3b). Both C>T and T>C substitutions are described as the predominant type of substitution in homoplasmic variants originating from germline transmission ^25,32^. The mutational signatures of heteroplasmic group C variants seems to be mainly shaped by the same mechanism, however, abundance of other substitution events suggest an additional origin such as EC culture expansion or tissue mosaicism of those variants. Consequently, group C mutations would, therefore, never have been subjected to the control and repair mechanisms acting during mtDNA transmission in germ cells.

Depending on the dynamic of mtDNA mutation progression during culture expansion, all group C variants in iPSCs were divided into four groups, namely enriched, stable, depleted, and fluctuating variants (Table 2, illustrated by the changes in blue shades). While enriched variants are defined as such that increased in AF over iPSC expansion culture more than 10 fold (average 76 fold; range 11-250 fold), stable variants did not change their AF during iPSC expansion culture (average fold change between passages 1.03; range 0.67-1.49; SD 0.25). Depleted variants were reduced in AF from the early or intermediate passage to the late one by more than 10 fold (average 64 fold; range 15-225 fold) (Table 2). The group of fluctuating variants incorporates basically variants that might be also added to one of the above mentioned groups, but as their fold change over culture only ranged from 2.7-8.8 fold (average 4.7; SD 2.2), we decided not to include them. A last group of variants (named “nd in p30-50”) shown in Table 2 comprises those iPSC-enriched variants that were found in iPSC clones that were only analyzed at early passage but not additionally in later ones.

Among the entirety of group C variants (AF > 0.02) all enriched variants were predicted to be neutral as well as 83% of the stable variants. The group of fluctuating variants or those not analyzed in high passages (nd in p30-50) contained 43%-50% putatively actionable variants (Fig. 2a). Most interestingly, the highest proportion of 88% of putatively actionable variants was observed among the depleted variants (Fig. 2a). Focusing only on group C variants with intermediate heteroplasmy levels (AF > 0.1, which equals an averaged heteroplasmy level of > 10%), yielded similar results, although the sample size of such was quite small (Fig. 2b). Thus, putatively actionable group C variants with intermediate heteroplasmy level were mostly depleted during culture, while neutral variants were unchanged in their heteroplasmy levels or became even enriched during culture expansion (Table 2). Fluctuating variants whose dynamics over culture was difficult to characterize, exhibited mostly only low AF. In general, the donor age did not affect the genetic stability or clearance mechanisms as in all groups (enriched, stable, depleted, and fluctuating) heteroplasmic group C variants are equally apportioned in neonatal (hUVEC) and aged (hSVEC) donors (Table 2).

**Fig. 2:**
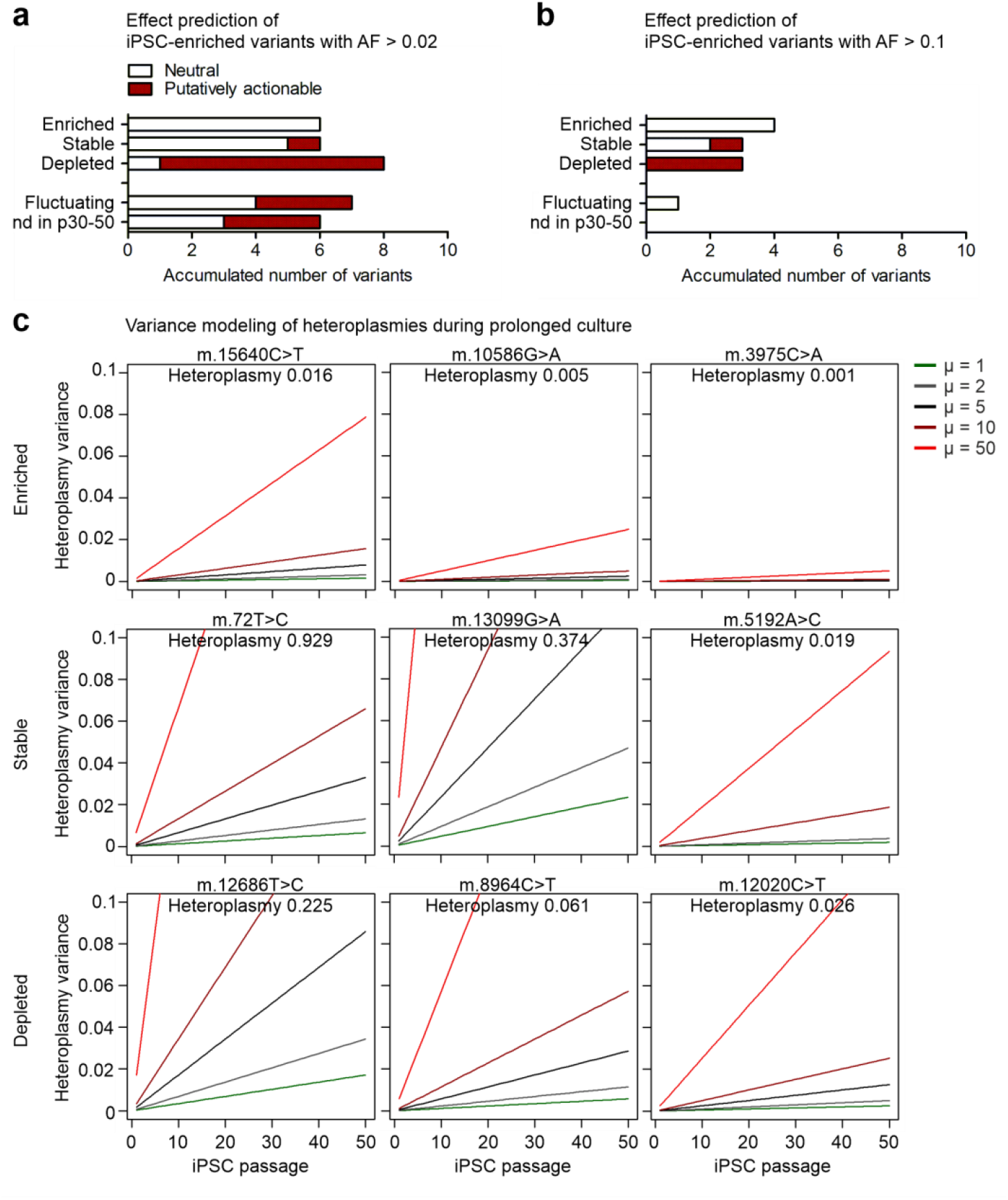
Prolonged culture expansion selects against putatively actionable mtDNA mutations in iPSCs. 26 iPSC clones at early passage (on average p6.5) and 7 of them additional at intermediate (p30), and at late passage (p50) were analyzed by mtDNA sequencing. Heteroplasmic variants (group C) were divided into the groups of enriched variants (defined as >10 fold increase in heteroplasmy level during culture), stable variants (with fold change ~1), depleted variants (decreased by > 10 fold), and fluctuating variants (change of heteroplasmy level did not allow confident classification into one of the above mentioned groups). The group of “nd in p30-50” comprises variants that were detected in iPSC clones that were only analyzed in low passages without exploring the dynamic of heteroplasmy. The ratio of neutral and putatively actionable iPSC-enriched variants within the group of enriched, stable, depleted, fluctuating, and “nd in p30-50” variants was determined. Effect prediction is based on a consensus of *in silico* prediction algorithms obtained from snpEff impact, CADD, Condel, and HmtVar. **a** Analysis including all heteroplasmic group C variants with minimum AF 0.02 at any passage of iPSC expansion culture. **b** Analysis including all group C variants with minimum AF 0.1 at any passage. **c** Modeling of heteroplasmy variance over passages, exemplarily for three variants with high, intermediate, and low heteroplasmy levels for each group. The model for genetic drift was adapted from Aryaman et al. ^24^ and allows prediction of variance of the heteroplasmy level and therefore, possible change of the heteroplasmy level over time. As neutral model, it predicts heteroplasmy variance only based on random gene drift, while selection forces can be incorporated by an additional coefficient. For the different heteroplasmy levels in iPSCs with N = 1000 mtDNA molecules, the heterplasmy variances in a neutral situation (green line) and with selection acting with various strength (μ = 2, 5, 10, and 50) are shown.

It is assumed that iPSCs are subjected to a genetic bottleneck when the mtDNA copy number per cell is reduced in the pluripotent state compared to somatic cells. It is still controversially discussed, however, whether random genetic drift during this bottleneck or selection processes are responsible for the observed changes in heteroplasmy levels during prolonged iPSC culture expansion ^21,24^. Notably, we did not observe any strong reduction in the mtDNA copy number in iPSCs compared to ECs of the parental cell populations (Fig. 1a). Furthermore, the mtDNA copy number per iPSCs stayed constant during prolonged iPSC expansion culture (Fig. 1b).

Here, we applied a neutral model of mtDNA genetic dynamics of Aryaman et al. ^24^ to understand how distribution of heteroplasmies evolves over time without selection. Therefore, this model was utilized as null hypothesis to assess whether random genetic drift by itself is sufficient to explain the observed changes in heteroplasmy levels, or whether additional selection factors play a role. The model of Aryaman et al. was developed for prediction of a possible change in heteroplasmy level of one mutated allele in a post-mitotic cell with mtDNA undergoing turnover. That means, degradation and replication of mtDNA occurs with the same extent over time allowing the ratio of mutated allele to fluctuate. Although our culture consists of mitotic iPSCs and requires frequent cell culture splitting, we applied this model here under the assumptions that i) iPSC experience symmetric division and equal segregation and expansion of all mtDNA molecules and ii) the effect of random gene drift during iPSC culture passaging is negligible. A more complex model predicting dynamics and selection in a unicellular life cycle with bottleneck and expansion in a multi-cellular organism ^23^ confirms that these assumptions prove valid for our population size (cells in culture before and after splitting). The model of Aryaman et al. includes, in addition, a fragmentation factor, which indicates the ratio of fused mitochondria. As unfused, punctuated mitochondria are a typical feature of iPSCs (Fig. S1), we excluded this factor in our modelling. The rate of turnover of mtDNA molecules and also selective mitophagy is set in this model by the factor *μ* (mitophagy rate). This factor is equivalent to a selection coefficient in other models ^23^.

Determination of mtDNA content has shown that the mtDNA copy number stayed constant over time during iPSC culture expansion (on average ~1000 copies / cell in 9 iPSC clones that were also analyzed in higher passages) (Fig. 1b). Thus, we simulated the possible heteroplasmy variance over time for three different heteroplasmic group C variants exemplarily for each group of enriched, stable and depleted variants, assuming cells contain 1000 mtDNA molecules and no selection is acting (Fig. 2c, green line). The variance in heteroplasmy level in cultured iPSCs increases with time according to random genetic drift. However, the modelling also indicates that expectable changes in heteroplasmy due to random genetic drift without computing an additional selection coefficient are very low even over the course of 50 iPSC passages (Fig. 2c, green line). Consequently, this result is consistent with the observation made in the group of stable group C variants which proved to be unchanged over the course of time (fold change on average 1.03, Table 2). However, this simulation without additional selection factor cannot explain the observed increase or decrease in heteroplasmy levels of the enriched or depleted variants. In contrast, incorporating a selection coefficient (included as μ factor) in the model can explain our observation (Fig. 2c). For depleted variants, a selection coefficient of μ = 10-50 (Fig. 2c, dark and light red lines) was, in general, sufficient to simulate the reduction of heteroplasmies. In contrast, to reach the change in heteroplasmy level observed by enriched variants, μ > 50 would even be needed in most of the cases. Taken together, the analysis of the mtDNA copy number and modelling revealed two aspects: i) the mtDNA copy number in iPSCs is high enough (no strong bottleneck in the pluripotent state) to prevent noteworthy effects by random genetic drift. ii) enrichment of neutral and clearance of putatively actionable heteroplasmic (group C) variants is likely caused by active selection during iPSC expansion culture.

### Dynamics of mtDNA copy number during differentiation does not change variant heteroplasmy levels

To investigate the development of heteroplasmy levels during differentiation and the consequences of mtDNA mutations for differentiated iPSC derivatives, the two iPSC clones harboring intermediate level of putatively actionable ND5 mutations (D#3 hUVEC C2 with 38% m.13099G>A and D#37 hSVEC C10 with 23% m.12686T>C) and their sister iPSC clones (D#3 hUVEC C1 and D#37 hSVEC C4) derived from the same parental cell population were differentiated into cardiomyocytes (CM). Monitoring of the mtDNA copy number during differentiation process, surprisingly, revealed a more complex dynamic than expected. Instead of a simple increase of mtDNA copies from iPSC to CMs with higher metabolic demands, the general dynamic followed a process through 3 phases (Fig. 3a). During the first phase (differentiation day (dd) 0 to dd1), starting with Wnt pathway activation by CHIR99021 treatment for 24h, mtDNA copy number per cell increased. After subsequent Wnt pathway inhibition by Wnt-C59 for 48h (dd1-dd3), mtDNA copy number per cell was reduced reaching its minimum around dd5 or dd6. After dd6 mtDNA copy number increased again, similar as described for cell differentiation during embryonic development ^40^, up to ~ dd10. The mtDNA copy number from dd10 onwards stayed unchanged with, on average, 1532 mtDNA copies per cell (Fig. 3a). Therefore, our results evidence that mtDNA copy number experienced two expansion phases and a reduction during the course of targeted CM differentiation. During reprogramming, such a bottleneck effect is suspected to exert influence on mtDNA segregation. However, analysis of heteroplasmy levels during the differentiation process by mtDNA sequencing at dd0, dd5, and dd15 of differentiation demonstrated that the heteroplasmy level of all variants stayed constant during differentiation with an average fold change of 1.00 (range 0.83-1.15; SD 0.08) (Fig 3b), a result supported by the observations of previous work ^14,21^. Therefore, a reduction in mtDNA copy number as assumed for a genetic bottleneck itself does not lead to change of average heteroplasmy level.

**Fig. 3:**
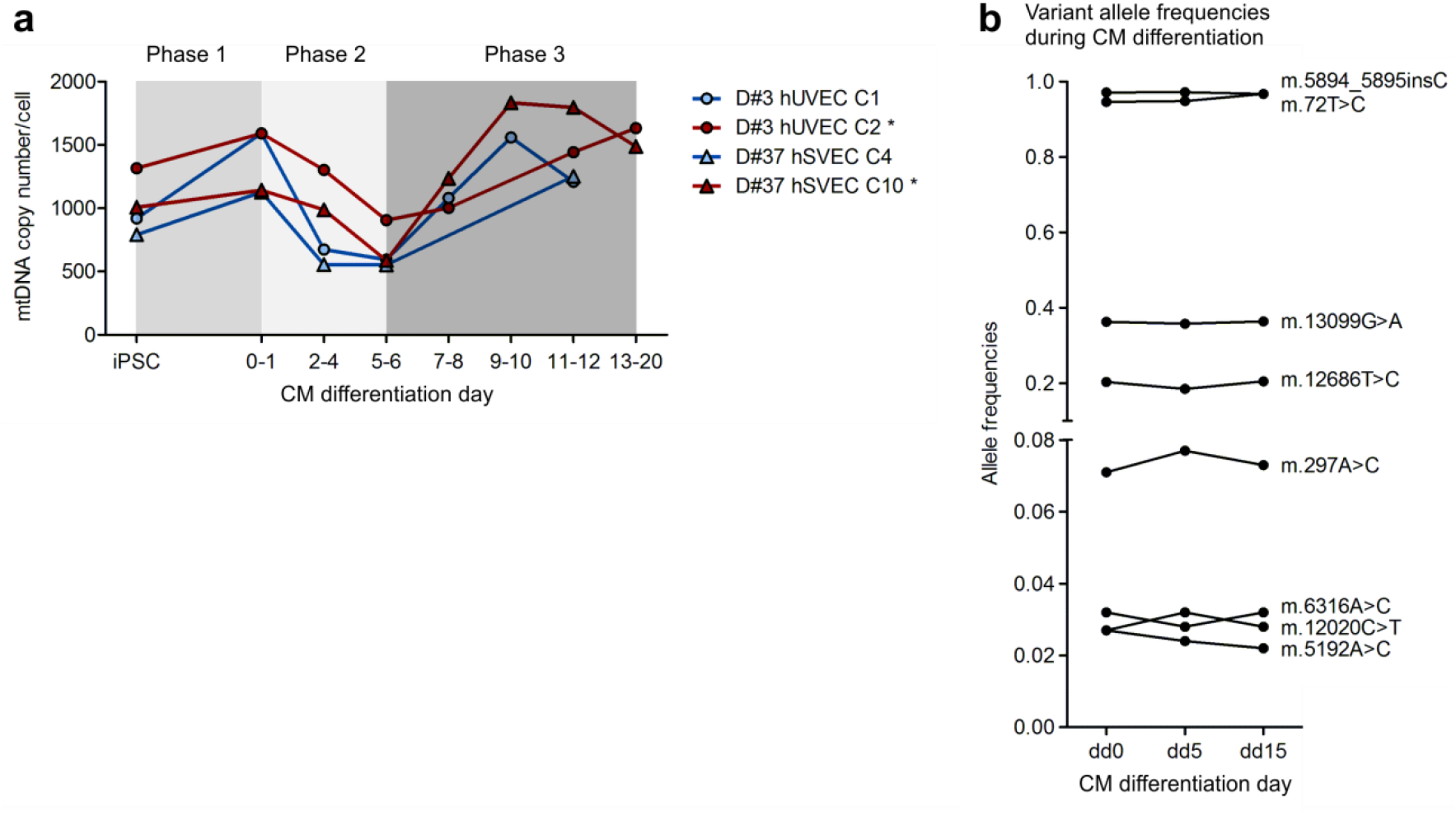
mtDNA content experiences reduction during cardiomyocyte differentiation without change in variant heteroplasmy levels. 2 iPSC clones (D#3 hUVEC C2 and D#37 hSVEC C10) harboring each a different putatively actionable mutation in ND5 gene (m.13099G>A and m.12686T>C, respectively) at intermediate heteroplasmy level and their isogenic sister iPSC clones were differentiated into cardiomyocytes (CMs). Mutated iPSC clones are marked with *. **a** Average mtDNA copy number per cell during CM differentiation. N = 1-2 differentiations. **b** At different time points during CM differentiation (differentiation day (dd) 0, dd5, and dd15), mtDNA of cells was sequenced. Plot displays the dynamic of heteroplasmy levels during CM differentiation of heteroplasmic variants with AF > 0.02. CM differentiation N = 4 clones with 1differentiation.

Moreover, mtDNA copy number dynamic was not influenced by presence of the ND5 mutations in iPSC clone D#3 hUVEC C2 or D#37 hSVEC 37 C10 (Fig. 3a, marked with *). Taking into account the relatively low variant allele frequencies of 0.38 (m.13099G>A) and 0,23 (m.12686T>C), it is, however, not unexpected that differentiation efficiency, final mtDNA copy number in CMs, and the phenotype of the mitochondria network were similar in the iPSC clones with and without the respective ND5 mutations (Fig. S4).

## Conclusion

Random genetic drift, positive selection or clearance of mutated mtDNA, are suspected to shape the mutational landscape of mtDNA in iPSCs, but to which extent and at what stage those partly contradicting processes might act is widely unknown. In any case, occurrence of mtDNA mutations that alter the functionality of iPSC-derivatives would exclude their clinical application. More detailed investigation of *de novo* mutagenesis, genetic drift, positive and negative selection processes during reprogramming, culture expansion and differentiation is therefore mandatory, as well as exploration of functional consequences of mitochondrial variants in iPSCs and their derivatives.

Here, we have analyzed mtDNA variants in iPSCs and their corresponding parental cell populations during reprogramming, prolonged culture, and differentiation. We neither observed a distinct bottleneck nor any selection for or against mtDNA variants during reprogramming. Overall, the number of mtDNA molecules per iPSCs was similar to the one in parental ECs. All mtDNA variants detected in iPSC clones have also been detected at different levels in the parental cells, although not in all cases with statistical confidence. Thus, we did not observe any evidence for substantial *de novo* mutagenesis, and inter-clonal differences in iPSCs apparently arose from the mitochondrial variant mosaicism within the parental cell population. During reprogramming, neutral and putatively actionable mtDNA variants were similarly transmitted from parental cells to iPSCs.

In contrast, during prolonged culture of iPSCs the heteroplasmy level of variants changed, while the mtDNA copy number per cell stayed uniform. Importantly, modelling the effect of random genetic drift on heteroplasmies revealed that a neutral model without selection cannot explain the observed changes. Therefore, heteroplasmies experience selection during the iPSCs stage. Interestingly, the selection acts pivotally against putatively actionable mutations, while neutral variants are accepted or even enriched. On a functional level, no difference in cultured undifferentiated cells between sister iPSC clones with and without ND5 mutation at intermediate heteroplasmy level derived from the same parental cell population was observed. Although heterogeneity among single iPSCs ^14^ might have led to elimination of damaged cells within the iPSC clone culture, this scenario is questionable as simultaneously other variants (stable group C variants) were maintained at a constant heteroplasmy level. It is more likely that small functional impact of mutations on mitochondria level led to selective mitophagy of damaged mitochondria and mtDNA via PINK1 and Parkin pathway, similarly as described for mtDNA integrity maintenance during germline transition ^29,34–36^. Hence, the clearance of mtDNA mutations is presumably rather executed by an intrinsic process on the intra-cellular level than on an inter-cellular level.

Importantly, our results demonstrate that during targeted CM differentiation the mtDNA copy number per cell is subjected to a distinct dynamic including an increment followed by a strong reduction to half of the mtDNA copy number, and lastly a two fold increase. Interestingly, this dynamic does not influence mtDNA average heteroplasmy levels at all so that the average heteroplasmy level from iPSCs remained unchanged in the differentiated cells. This observation leads to two conclusions: i) the heteroplasmy level in iPSCs dictates the heteroplasmy level of the iPSC-derived CMs. ii) the observed dynamics of the mtDNA copy number during the course of the CM differentiation seems to be not yet strong enough to cause unequal mtDNA segregation.

Altogether, our results suggest that the bottleneck effect might have been overestimated, in context of the mtDNA genetic pool. Our findings point towards a scenario in which intra-cellular positive and negative selection of mtDNA molecules mainly shapes the mutational landscape of the mitochondrial genome in iPSCs while other mechanisms such as random genetic drift and inter-cellular selection (clonal eliteness) exert a minor impact. Above all, similar to female germline development and early embryogenesis ^16,29,33^, this selection purges putatively actionable mtDNA mutations averting the mutational ratchet in iPSCs.

## Methods

### Reprogramming and iPSC culture

Derivation and culture of endothelial cells, virus production, retroviral reprogramming, and iPSC characterization was performed essentially as previously described ^41^. In short, endothelial cells (ECs) were isolated from umbilical vein (hUVEC) from healthy newborns and saphenous vein (hSVEC) from aged patients (64-88 years) that underwent coronary bypass surgery. Human material was collected after approval by the local Ethics Committee and following the donor’s or the newborn’s parental written informed consent. ECs were cultivated in Endothelial Growth Medium (EGM-2) (Lonza) and 2*10^5^ cells of early passages (mean p4.5; range p3-7) were reprogrammed by ectopic expression of Oct4, Sox2, Nanog, and Lin28 (lentiviral transduction, monocistronic factors, multiplicity of infection (MOI) 20) with exception of clone 5 of donor D#40 hSVEC (D#40 hSVEC C5) which was reprogrammed via ectopic expression of Oct4, Sox2, Klf4, and c-Myc (lentiviral transduction, monocistronic factors, MOI 1). In total, 26 EC-derived iPSC clones were derived from 10 donors. Thereby, 3 clones were generated for each donor D#1 hUVEC-D#5 hUVEC, D#31 hSVEC, D#37 hSVEC, and D#38 hSVEC, while only 1 clone was selected for the donors D#6 hUVEC and D#40 hSVEC. iPSC clones were cultivated as colonies on mouse embryonic fibroblasts (MEF) in Knock-out Dulbecco’s Modified Eagle’s medium (DMEM) (Gibco) supplemented with 20% KnockOut serum replacement, 1% Non-essential Amino acids, 1mM L-Glutamine, 0.1mM β-Mercaptoethanol (all obtained from Life Technologies), and 10ng/ml bFGF (Institute for Technical Chemistry, Leibniz University Hannover, Germany). Before execution of any analysis, iPSCs were transferred to monolayer culture on Geltrex (Thermo Fisher Scientific) in in-house Essential 8 (E8) medium (DMEM Nutrient Mixture F-12 (Gibco) supplemented with 543mg/L NaHCO_3_, 2ng/ml TGFβ (PeproTech), 10.7μg/ml human recombinant transferrin, 14μh/l sodium selenite, 20μg/l insulin, 64μg/l ascorbic acid 2-phosphate (all obtained from Sigma-Aldrich), and 100ng/ml bFGF) and cultured for generally 3-5 passages.

### Total DNA extraction and determination of mitochondrial DNA copy number using quantitative real-time PCR (qRT-PCR)

Total DNA was isolated using QIAamp DNA Blood mini kit (Qiagen). A quantitative real-time PCR (qRT-PCR) based analysis system was developed which quantifies mitochondrial DNA (mtDNA) copy number relative to genomic DNA (gDNA) copy number. In contrast to most other studies which use only one primer pair, our system measures number of mtDNA and gDNA copies by each three primer pairs. Sequences for two primer pairs targeting mtDNA regions were assimilated from previous studies ^42,43^. The third primer pair for mtDNA and those for gDNA quantification were designed employing Primer3 (v0.4.0). During primer design and selection, we made sure that primers did not target loci within the common 4700bp mtDNA deletion site or genomic regions with high CNVs or mutation occurrence. Loci were manually inspected using variation data provided by Ensembl (release 90, August 2017) based at European Bioinformatics Institute (EMBL-EBI) (Hunt, Ensembl variation resources, Database 2018). The primer list is presented in Table S4. qRT-PCR was performed on 25ng total DNA per reaction in duplicates on Mastercycler ep realplex2 (Eppendorf) and with Absolute QPCR SYBR Green Mix (Thermo Scientific). Size of amplicons and absence of nonspecific byproducts were confirmed via melting curve analysis using realplex software (Eppendorf). Quantification was obtained by both calculation of ddCt values and standard curve comparison. Both quantification approaches always resulted in comparable results. mtDNA copy number per cell was calculated as average of mtDNA copy number measured by the three primer pairs relative to average gDNA copy number/2.

### Enriched mtDNA extraction and mtDNA sequencing

The workflow for mtDNA sequencing and sequencing data processing is displayed in Fig. S5a. Enriched mtDNA was extracted via QIAprep Spin Miniprep kit (Qiagen) from early passages of iPSC clones (mean p6.5; range p3-10) and from late passages (p30 and p50) of 7 clones. Furthermore, enriched mtDNA was isolated from the corresponding parental cell populations at a similar passage as subjected to reprogramming (mean p4.5; range p3-7). Isolation was essentially performed according to manufacturer’s recommendation. Enriched mtDNA isolation yielded, on average, 12 fold increase of mtDNA compared to total DNA isolation approach (Fig. S5b). Entire mtDNA was then amplified using 2 primer pairs (F2480A/R10858A in ratio 3:1 and F10653B/R2688B in ratio 1:1) (Table S4) generating two overlapping PCR products of 8379bp and 8605bp fragment size^44^. PCR reactions were performed with Herculase II Fusion DNA Polymerase (Agilent) in technical replicates with 50ng enriched mtDNA as template per reaction. Altogether, the input of enriched mtDNA accounted for 200ng which equals to ~ 5*10^8^ mtDNA molecules (3.2*10^4^ cells (theoretical input) * 12 (average fold enrichment) * 1300 (average mtDNA copy number per cell)), isolated from, on average, 5*10^6^ iPSCs or 2.5*10^6^ parental cells. PCR conditions were: 94°C for 5min as initiation step, then 23 cycles of 94°C for 40s, 54°C for 20s, and 54°C for 4.15min, followed by termination step of 72°C for 3min. Therefore, our protocol reduced PCR amplification cycle to 23 compared to 30 in most other published approaches of mtDNA amplification for sequencing purposes. mtDNA PCR amplification resulted in an enrichment of mtDNA of, on average, ~15000 fold over normal cellular content (Fig. S5b). Each amplicon was individually inspected by gel electrophoresis and amplicons from the same sample were then pooled, purified applying AMPure XP beads (Beckman-Coulter) in 0.6x ratio, sheared by focused-ultra sonication (Covaris), and quantified using a Qubit fluorimeter (Invitrogen). Sheared fragments were purified with AMPure XP beads (0.9x ratio) and 150ng of each sample were fed to library preparation with NEBNext Ultra II DNA Library Preparation kit for Illumina (New England BioLabs) according to manufacturer’s instruction. The final concentration, fragment distribution (mean fragment length 600bp; range 500-700bp), and quality were assessed on Qubit fluorimeter and Agilent Bioanalyzer. The library was sequenced as paired-end 250bp reads using MiSeq Reagent Kit v2 (Illumina).

### Read alignment, variant calling and refinement

Post-run fastq files were imported into Galaxy (v17.05) ^45^ instance of the RCU Genomics, Hannover Medical School, Germany for read trimming, quality assessment, alignment, and variant calling. Quality and adapter trimming of reads was performed by Trim Galore (v0.4.3; Cutadapt v1.14) including removal of adapter, 10bp from the 3’ end of read 1, 30bp from the 3’ end of read 2, as well as low-quality ends from reads (Phred < 28). Reads were aligned to the human genome GRCh38 chromosome MT (the Cambridge Reference sequence (rCRS; NC_012920)) using BWA-MEM (Galaxy Version 0.7.17.1) 46. Quality of trimmed and aligned reads were inspected employing FastQC Read Quality reports (v.0.11.6; Galaxy wrapper version v0.71) and Qualimap Multi-sample BAM QC analysis (v.2.2.1). Reads had an average length of 215bp, with Phred score > 28 over whole read length. 98% of all reads mapped to mitochondrial reference genome with equal quality across the reference genome (59.8 mean mapping quality). The coverage was 19000 on average.

Variants were called utilizing FreeBayes (v1.0.2; Galaxy implementation v1.0.2.29-3) ^47,48^ with filters set to ploidy 10, minimum coverage 100, and min AF (alternative frequency) 0.02. Min AF of 0.02 as filter criterion was established by manually reviewing called variants within their read context. A receiver operating characteristic (ROC) curve with AF as discrimination threshold (Fig. S5c) showed that min AF 0.02 retrieve very high specificity while maintaining most true positive variants. False variants with min AF 0.02 comprised mainly remaining adapter sequences that evaded the trimming process and presumable sequencing errors of C-stretch or C-rich genomic regions. To exclude the few remaining false positive variants, remaining primer sequences were manually removed and variants that were detectable in different donors (with min AF 0.005) but not described as polymorphisms were excluded as probable false variant calls.

Furthermore, to investigate presence of a choice of variants at very low AF (in general, < 0.02) in iPSC clones or assess their pre-existence in parental cell population, in total 128 variants at 38 different genetic regions were examined individually within their genetic context. Therefore, sequencing reads were inspected directly from fastq raw files. Both reads of the paired end sequencing were investigated and the number of reads with the variant enclosed by a 11bp long sequence ranging from position −5bp to +5bp was counted as well as the number of reads with the reference sequence. For calculation of average error for each genetic location, the numbers of reads that comprised not the variant or reference nucleotide were determined as well. In addition, the 11bp long sequences directly upstream and downstream of the variant were analyzed in the same manner. The mean error rate and detection limit was calculated individual for every variant and presence of variant was confirmed with p-value 0.05 against background of errors (Fig. S5c, Table S2a). Frequencies of INDELs were determined by extracting and counting reads for both INDEL and reference sequence embedded by an 11bp sequence, directly from fastq files (Table S2a). Overall, the average coverage of the variant regions was 22000 and detection limit was 1 read in ~ 340 (SD 450) which equals to a detection sensitivity down to AF 0.003, on average.

### Variant annotation and prediction of functional consequences

Variant annotation was performed using the web interface of Ensembl Variant effect predictor (VEP) (v95) (assembly GRCh38.p12) ^49^. Haplogroups and corresponding haplotype variants were identified via HaploGrep algorithm ^50^. MITOMAP (A human mitochondrial genome database, r103) was utilized to retrieve population frequencies (NCBI GenBank frequency (GB)) of variants. Variants with GB frequency ≥ 0.02 were here considered to be polymorphic.

Prediction of functional consequences was obtained from Ensembl VEP and HmtVar ^51^. The functional consequence of a variant was classified by a consensus based on the in silico predictions of snpEff impact prediction, Condel, CADD (Ensembl modules), and HmtVar prediction. If a variant had a harmful designation by snpEff (high impact) or at least by two of the other algorithms (Condel=deleterious, CADD phred > 20, HmtVar=pathogenic), provided prediction tools returned a result, it was considered as putatively actionable.

### Modeling heteroplasmy variance over time

A model by Aryaman et al. (2019) was adapted for modeling heteroplasmy variance over time ^24^.

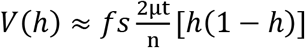

Where *f_s_* is a factor for fragmentation which indicates the ratio of fused mitochondria, *μ* is a factor for mitophagy rate, and *h* the heteroplasmy level. As iPSCs are featured by unfused, punctuated mitochondria, *f_s_* was excluded in our modeling. *n* represents the mtDNA copy number/cell. Determination of the mtDNA copy number in iPSCs (*n*) in p6, p30, and p50 showed that *n* stayed constant over time in culture and was, on average, 1000. Heteroplasmy variance (*V(h)*) was calculated for different heteroplasmy levels with different μ to assess effect of genetic drift (without selection, *μ* = 1) and selection (*μ* = 2, 5, 10, and 50).

### Differentiation of cardiomyocytes (CMs)

For embryoid body formation, (differentiation day (dd) −3) 1*10^6^ undifferentiated cells/well were aggregated in 3ml 10μM Rho-Kinase inhibitor Y-27632 (RI) containing E8 medium on low attachment 6 well (Greiner Bio-one GmbH) plates on an orbital shaker (70rpm; Infors GmbH). Differentiation was started (dd0) using 5μM CHIR99021 (LU Hannover) in CDM3 ^52,53^ to activate the WNT pathway. Exactly 24h later it was followed by a subsequent inhibition of the WNT pathway (dd1), where CDM3 medium including 2μM Wnt-C59 (LU Hannover) was added for 48h. On dd3 and dd5 fresh CDM3 medium was added. From dd7 on, the medium was changed to basic serum free medium (bSF; DMEM with 1% non-essential amino acids, 5.6mg/l transferrin, 37.2μg/l sodium-selenit, 1mM L-glutamine, and 10μg/ml insulin) and was changed every other day. The differentiation efficiency was addressed on dd14 by flow cytometry for the cardiac markers, cardiac troponin T (cTnT), alpha sarcomeric actinin (αSA), and pan-myosin heavy chain (MF20) as described earlier ^53^ (Table S5).

### Live-cell staining and flow cytometric analysis

Live-cell staining was performed of adherent cells with MitoTracker, ROS Brite 670, and the mitochondria-specific dye tetramethylrhodamine methyl ester (TMRM, 25nM; Thermo Fisher Scientific) diluted in Hanks solution (Gibco) with 20mM Hepes (Sigma-Aldrich) (HHBS) according to manufacturer’s recommendation for 45min. List of used antibodies and dyes is provided in Table S5. Subsequently, cells were dissociated by Versene (Gibco) or Typsin/EDTA treatment for iPSCs or ECs, respectively, resuspended in HHBS buffer, filtered, and analyzed by flow cytometry. Flow cytometry was executed on MACSQuant analyzer (Miltenyi Biotec) and data were analyzed with FlowJo (v10).

### Immunofluorescence staining and microscopic analysis

ECs, iPSCs, and CMs were seeded on cover slides coated with 1% gelatin, Geltrex, or 50μg/ml fibronectin in 0.02% gelatin, respectively. After a culture period of 1-2 days for ECs and iPSCs and 3 days for CMs, cells were fixed with 4% Paraformaldehyde (PFA) for 20min at 4°C, permeabilized with 0.1% Triton X 100 (Sigma-Aldrich) in PBS for 5min at room temperature, and unspecific binding sites were blocked with solution of 5% donkey serum (Chemikon) and 0.25% Triton X 100 in tris-buffered saline (TBS) for 30min at 4°C. Incubation with monoclonal anti-ATPIF1 (ATP synthase subunit IF1) antibody diluted in staining buffer (PBS with 1% bovine serum albumin (BSA) (Sigma-Aldrich)) was performed overnight at 4°C. List of used antibodies is provided in Table S5. Subsequently, secondary antibody staining was performed for 1h at 4°C and DAPI nuclear staining for 1min. Cells were analyzed with Axioserver A1 fluorescent microscope (Zeiss) and Axiovision (v4.71) software.

### Measurement of glucose consumption and lactate production rate

iPSCs were cultured as monolayer in house-made E8 medium. Day 1 after cell seeding medium was exchanged and at the two subsequent days, lactate and glucose concentrations from cell-free supernatant were measured employing Biosen C-Line glucose and lactate analyzer (EKF Diagnostics). Each day of analysis cell number were also counted with Neubauer chamber. Technical triplicates initiated from the same starting cell population were analyzed for each sample and time point. Yield coefficient of lactate from glucose was calculated as previously described ^54^.

### Statistical analyses

GraphPad Prism (v6.07) or RStudio (v1.1.463; R v3.5.1) software were employed for statistical analysis and visualization. Results are presented as mean and SD. D’Agostino and Pearson omnibus normality test were executed to check if samples were normally distributed. Subsequently unpaired two-tailed t-test and nonparametric two-tailed Mann Whitney test were employed as appropriate to test for significant differences between groups.

Average error of the amplicon sequencing was calculated for every variant as the mean number of reads with non-reference and non-variant nucleotide at variant location or in the middle of analyzed upstream and downstream sequence (in general, N = 8). Distribution of errors for each variant was evaluated via the D’Agostino and Pearson omnibus normality test (where applicable, else the Shapiro-Wilk normality test was used). One sample one-tailed t-test and Wilcoxon one-tailed signed-rank test were performed, respectively, to compare number of reads with variant as hypothetical value against background of errors. The null hypothesis, assuming number of reads with variant is within local error range, was rejected with p values of 0.05.

## Code availability

Sequencing data analysis was mainly performed on the Galaxy (v17.05) ^45^ instance of the RCU Genomics, Hannover Medical School, Germany. The workflow is available at Supplementary Data 2.

## Data availability

Data are stored on the HPC-seq clusterserver at Hannover Medical School, Germany and available upon request.

## Acknowledgements

Many thanks to K. Osetek for isolation of hUVEC and hSVEC, and their reprogramming into iPSCs, to P. Stiefel for several HSVEC isolations, and to T. Scheper for providing bFGF. We are also thankful to S. Rojas, T. Goecke and other surgical colleagues, who provided human tissue samples. We thank J. Thomson for providing the reprogramming plasmids (SIN-EF2-Lin-Pur, pSIN-EF2-Nanog-Pur, pSIN-EF2-Oct4-Pur, pSIN-EF2-Sox2-Pur) via Addgene and D. Trono for providing the lentiviral packaging and transfer plasmids psPAX2 and pMD2.G. Many thanks as well to A. Haase and S. Merkert for a critical reading of the manuscript.

## Author contributions

M.K. contributed to designing and coordinating the study, performed, experiments, collected and analyzed data, and wrote the manuscript. C.D. and L.W. performed sequencing and contributed to analyzing bioinformatics data. T.K. performed experiments. M.D. provided technical assistance in fragment library construction and sequencing. J.G., K.M., and M.Si provided technical assistance in performing experiments. M.Sz, I.G. and A.M. coordinated and performed cardiomyocyte differentiation and analysis. U.M. designed and coordinated the study.

## Supplemental information

**Fig. S1:**
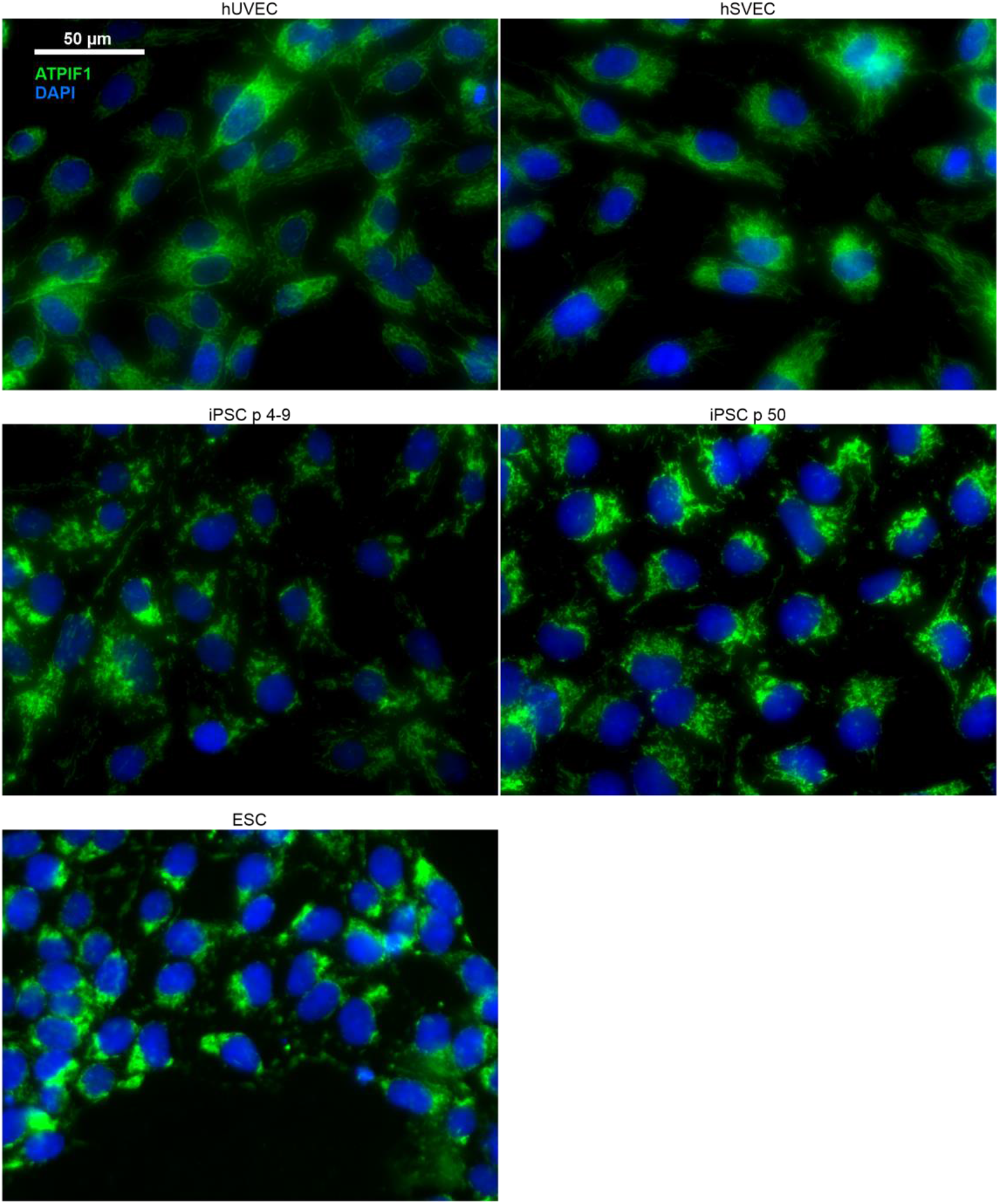
Mitochondrial network in ECs, iPSCs, and ESCs. Adherent ECs (umbilical vein (hUVEC) and saphenous vein (hSVEC)), iPSCs of early and late passage, and ESCs were seeded on gelatine or Geltrex coated glass cover slides, cultured for 2-3 days in EMG2 or E8 medium, fixed with PFA, stained with anti-ATPIF1 (ATP synthase subunit IF1) antibody and DAPI nuclear staining, and analyzed by fluorescence microscopy. Pictures of different representative donors and clones are shown. Mitochondria are displayed in green and nuclei staining in blue. Scale bar 50μm.

**Fig. S2:**
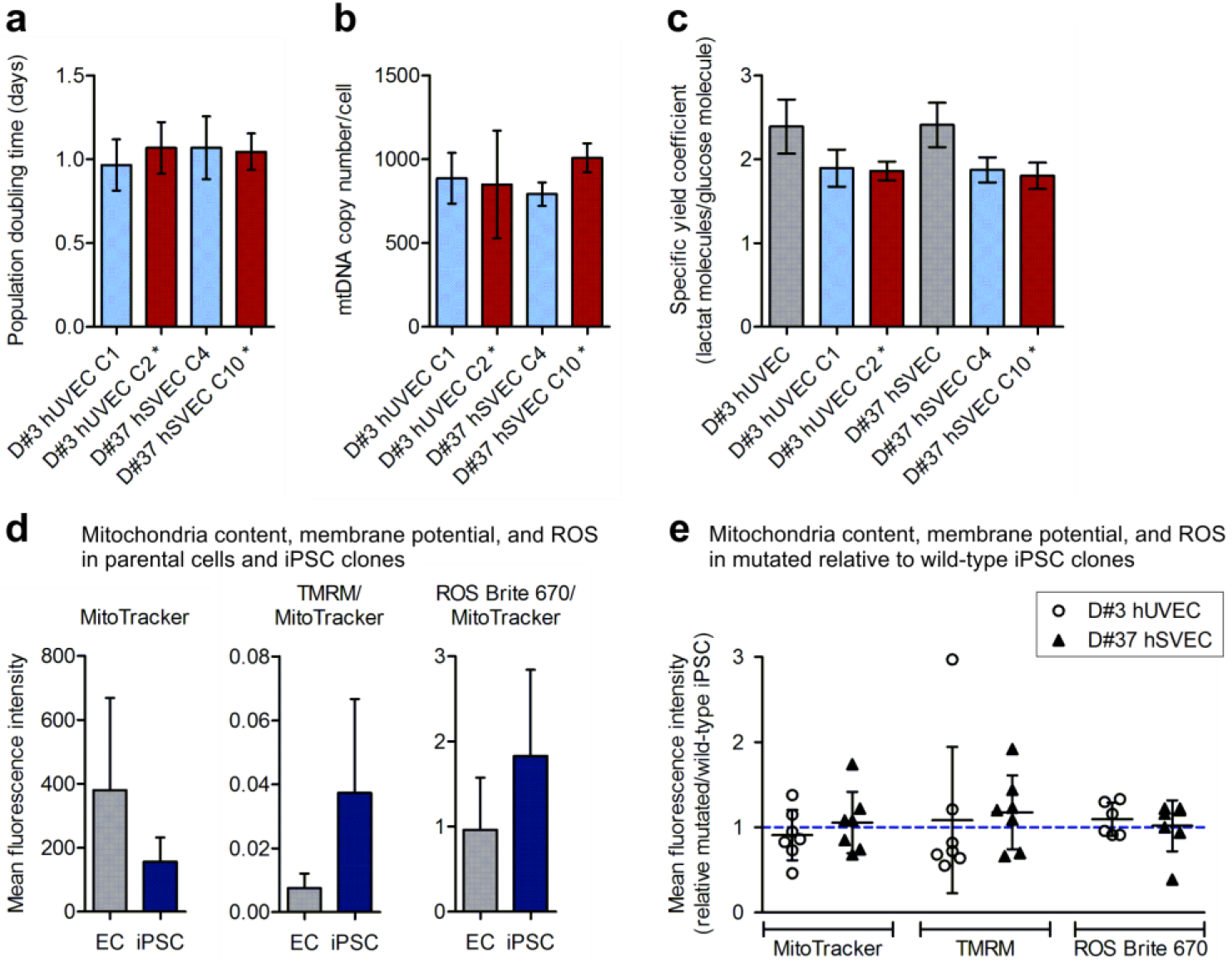
No detectable effect of specific ND5 mutations on the functionality of undifferentiated iPSCs. iPSC clones with putatively actionable ND5 mutations at an intermediate heteroplasmy level (D#37 hSVEC C10 with m.12686T>C at heteroplasmy level of ~20% and D#3 hUVEC C2 with m.13099G>A and ~36% heteroplasmy) and their sister iPSC clones without ND5 mutations derived from the same parental cell population were analyzed. Mutated iPSC clones are marked with *. **a** Population doubling calculated as time in days needed for doubling of cell number measured by cell counting. Cell numbers were determined over 8-12 passages with 3 technical replicates. Mean with SD. **b** mtDNA copy number per iPSCs. Mean with SD. N = 2-4. **c** Yield coefficient of lactate from glucose. Mean with SD. N = 2-4 with 2-3 technical replicates. **d-e** Determination of mitochondrial content (MitoTracker), membrane potential (TMRM), and reactive oxygen species (ROS Brite 670) per cell via live cell staining and flow cytometric analysis. **d** Quantification calculated as geometric mean of fluorescence intensity of staining-positive populations. Mean with SD. EC N = 4; iPSC N = 8 including 4 measurements of mutated and 4 of wild-type clones. **e** Comparison of metabolic features of mutated against wild-type iPSC clones. Plot displays the deviation of values measured in mutated iPSC clones from the values in their wild-type counterparts quantified by geometric mean of fluorescence intensity. Mean with SD. N = 7.

**Fig. S3:**
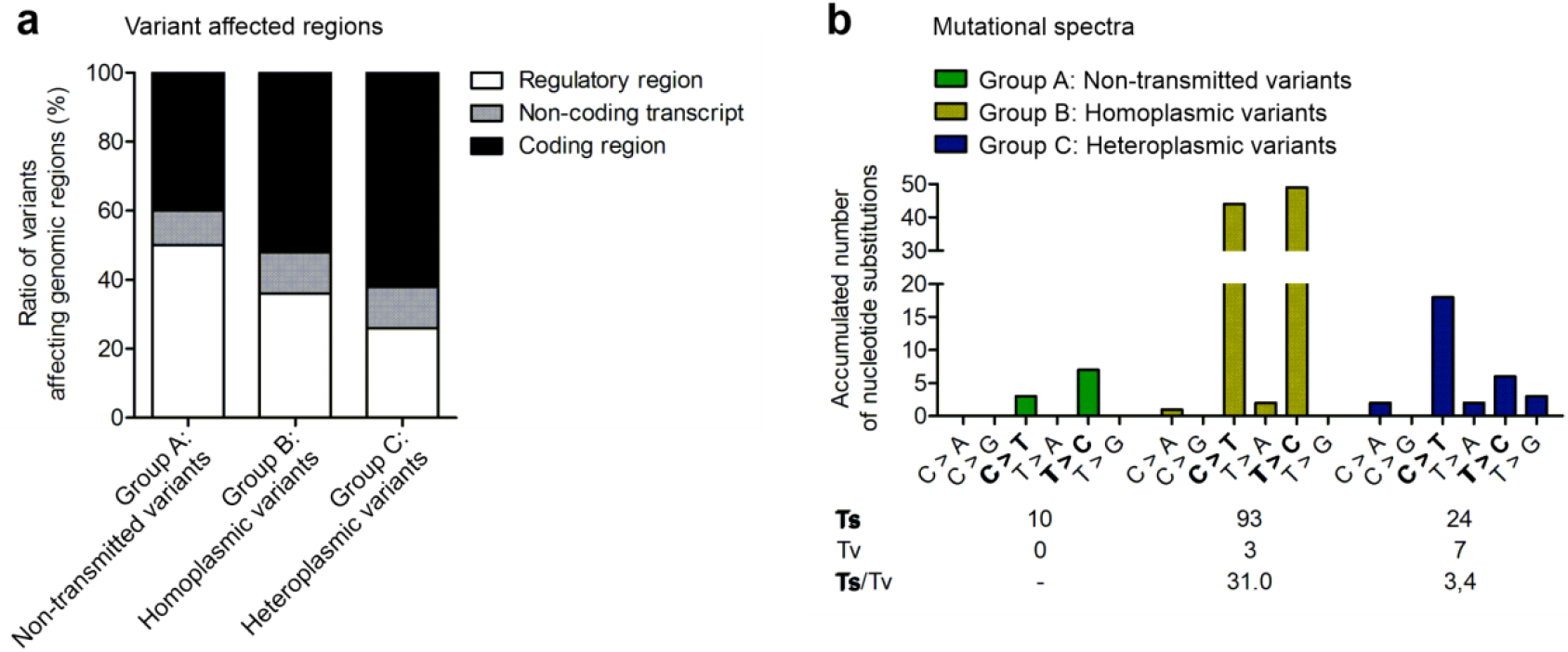
Transmitted heteroplasmic group C variants affect coding regions more frequently and are characterized by a higher transversion rate. 26 iPSC clones at an early passage, with 7 of these additionally at later passages, as well as the corresponding parental cell population of the total 10 donors, were subjected to mtDNA sequencing. All variants were classified into three groups. Variants of group A, non-transmitted variants, were detected only in parental cell populations. Group B variants were homoplasmic in parental cell population and iPSCs derived thereof. Group C, heteroplasmic variants, were detected in iPSC clones with AF > 0.02 at any passage but were less frequent in the corresponding parental cell population. **a** Total number of non-transmitted, homoplasmic, and heteroplasmic variants that affected regulatory, non-coding transcript, or coding regions. **b** Mutational spectrum of non-transmitted, homoplasmic, and heteroplasmic variants. Ts: transition; Tv: transversion. Transitions are marked in bold.

**Fig. S4:**
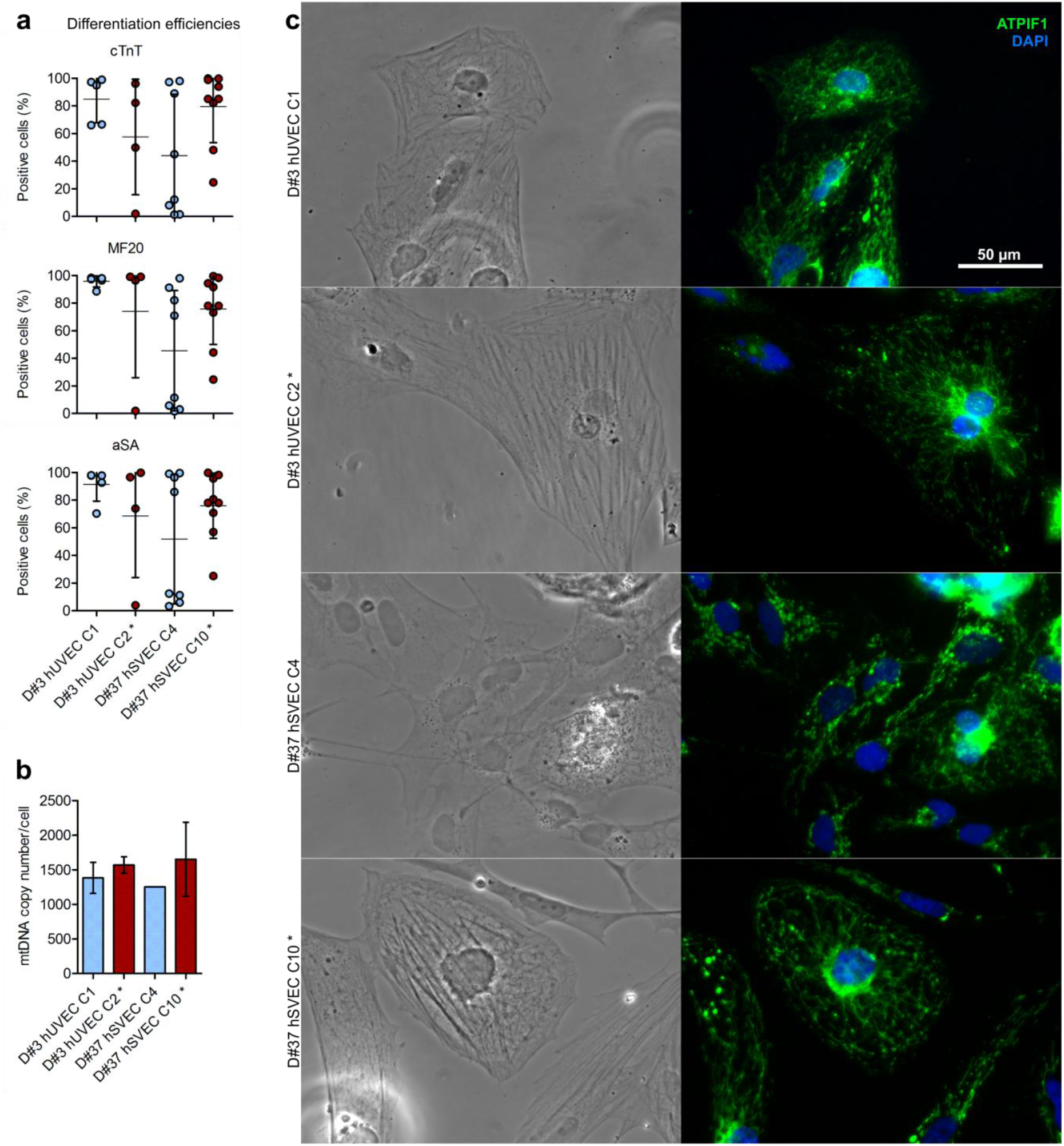
Intermediate heteroplasmy level of putatively actionable ND5 mutation does not alter differentiation potential of iPSCs into CMs. 2 mutated iPSC clones (marked with *; D#3 hUVEC C2 and D#37 hSVEC C10) each harbor a different putatively actionable mutation in ND5 gene at intermediate heteroplasmy level and their isogenic sister iPSC clones were differentiated into CMs. **a** Differentiation efficiency quantified by positive staining for cTnT (Troponin-T), MF20 (MYH1E), and aSA (Sarcomeric α-actinin) and flow cytometry analysis at differentiation day (dd) 14. Mean with SD. N = 4-9 differentiations. **b** mtDNA copy number per CM determined at dd10-20. Mean with SD. N = 1-2 differentiations. **c** CM at dd15 were seeded on fibronectin/gelatin (50μg/ml in 0.02% gelatin) coated glass cover slides, cultured for 3 days in differentiation medium, fixed with PFA, stained with anti-ATPIF1 (ATP synthase subunit IF1) antibody (green) and DAPI nuclear staining (blue), and analyzed by fluorescence microscopy. Scale bar 50μm.

**Fig. S5:**
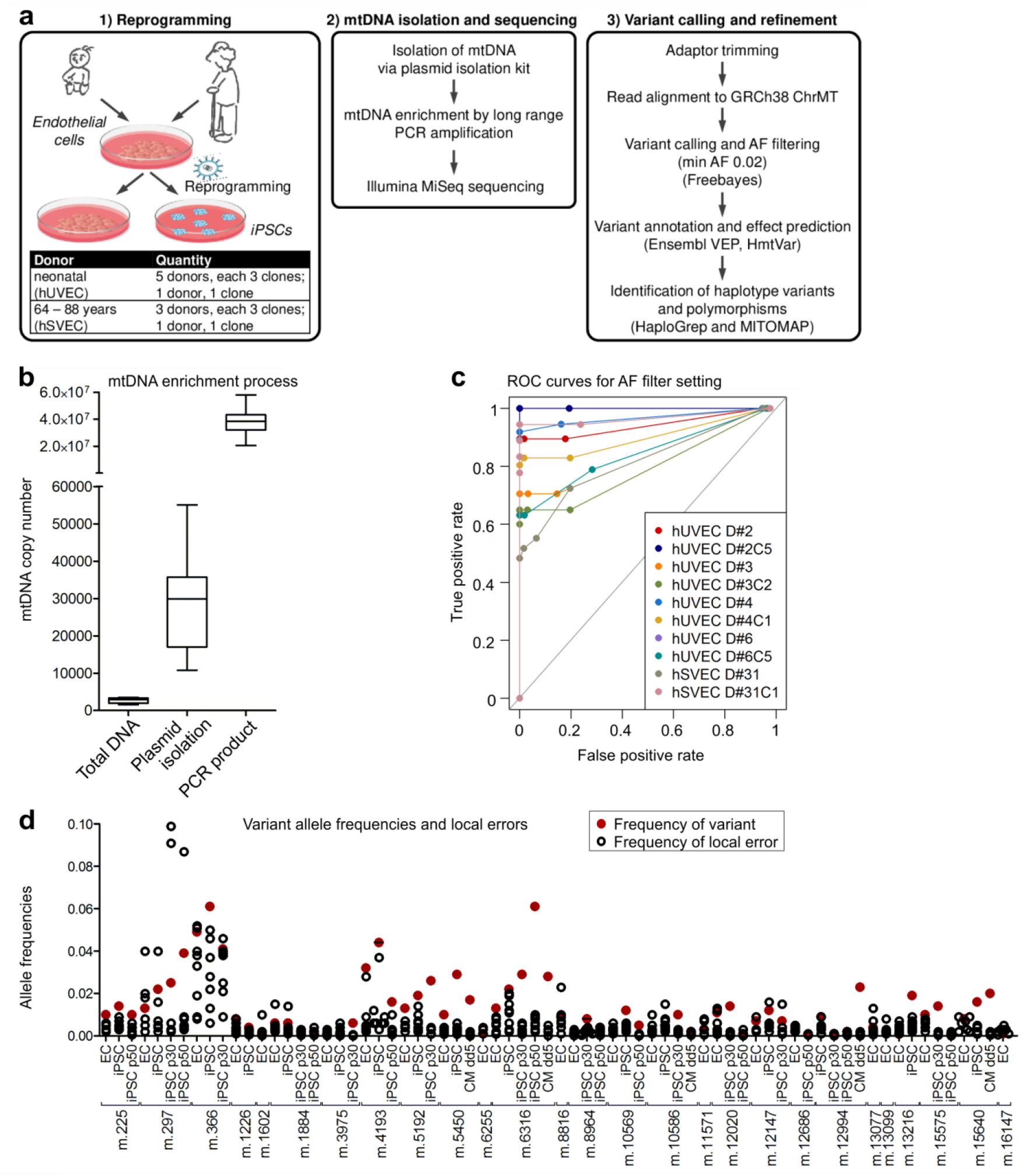
Establishment of our mtDNA sequencing approach. **a** Workflow including cell reprogramming, sequencing and sequencing data analysis. **b** mtDNA enrichment by plasmid isolation and long range PCR amplification for mtDNA sequencing. mtDNA copy number is measured relative to gDNA copy number after total DNA isolation, which represents normal cellular proportion of mtDNA and gDNA, after plasmid isolation, and after additional long range PCR amplification of mtDNA. **c** Receiver operating characteristic (ROC) curve with AF as discrimination threshold (AF 0.005, AF 0.01, AF 0.02, AF 0.04, AF 0.06, AF 0.08, AF 0.1, AF 0.5, AF 1). **d** A choice of variants with very low variant allele frequencies in iPSC clones and/or parental cells were additionally investigated individually within their genetic context by inspecting and counting sequencing reads directly. The plot shows the AF of single nucleotide variants (SNVs), calculated from the number of reads with variant, in red against the background of reads with the non-reference and invariant nucleotide exemplarily for 27 loci.

**Table S4:**
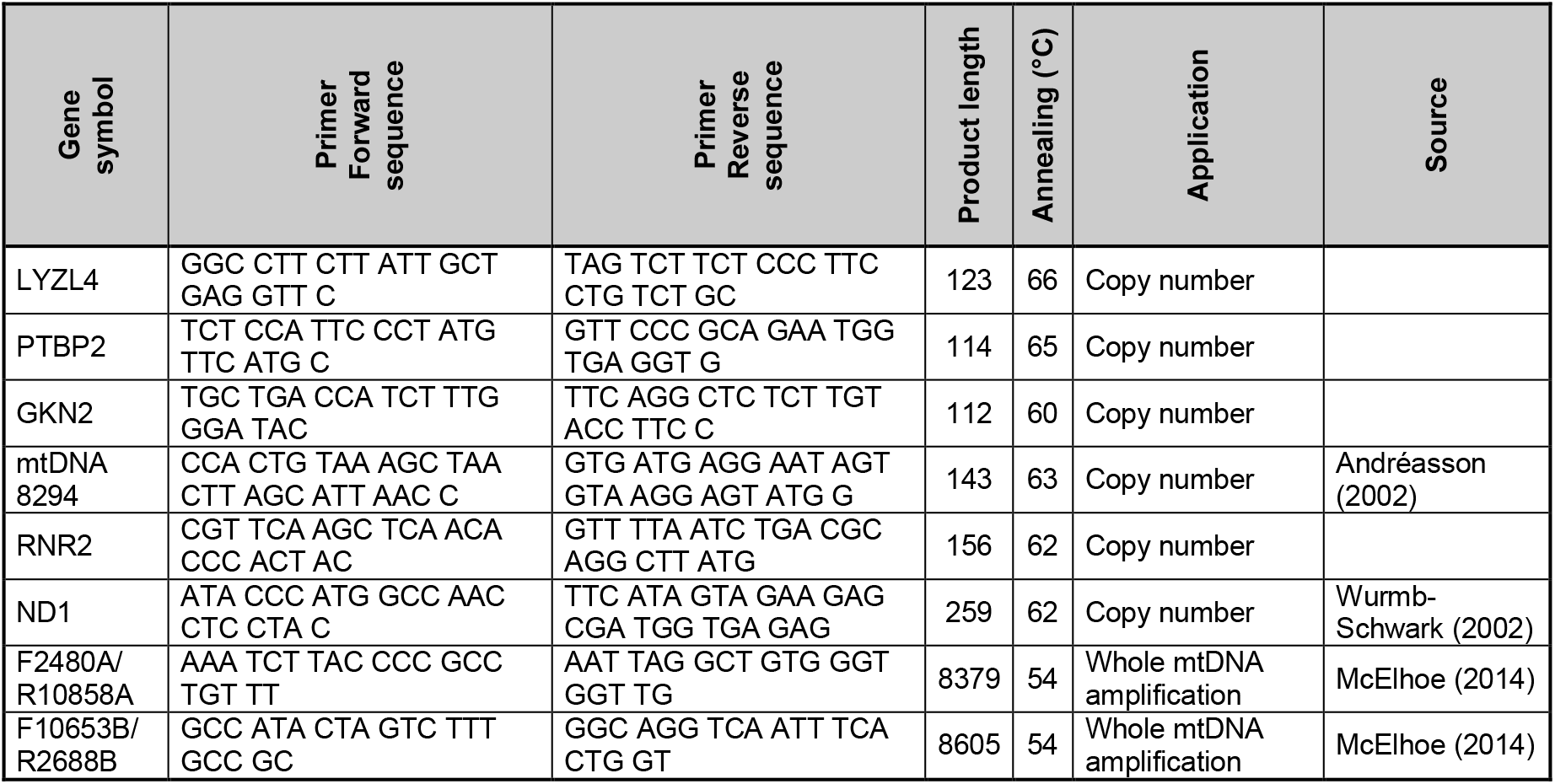
Primer list. Primer sequences for mtDNA copy number determination by qRT-PCR and whole mtDNA amplification for mtDNA sequencing.

**Table S5:**
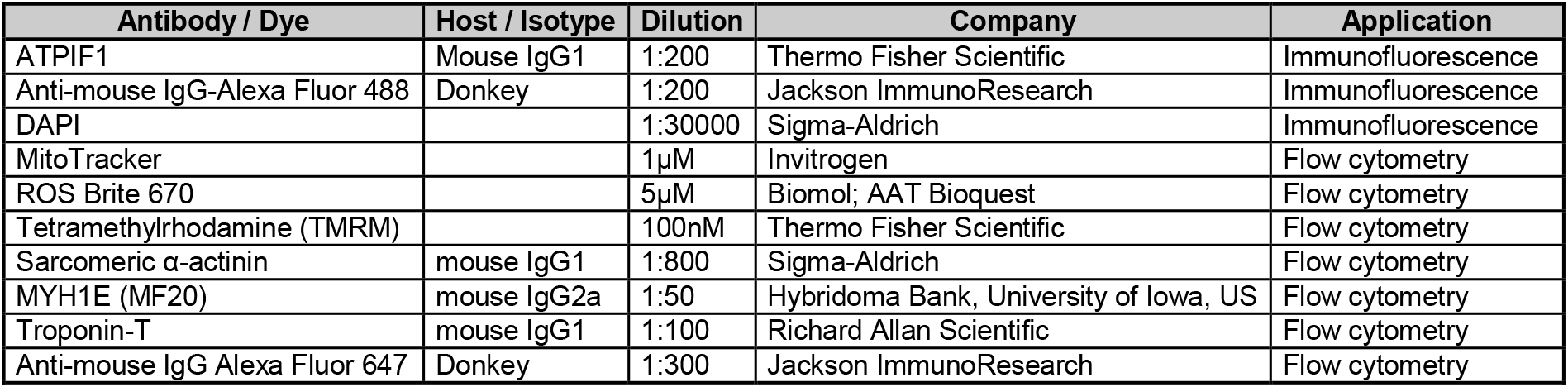
Antibody and dye list.

| Antibody / Dye | Host / Isotype | Dilution | Company | Application |
| --- | --- | --- | --- | --- |
| ATPIF1 | Mouse IgG1 | 1:200 | Thermo Fisher Scientific | Immunofluorescence |
| Anti-mouse IgG-Alexa Fluor 488 | Donkey | 1:200 | Jackson ImmunoResearch | Immunofluorescence |
| DAPI |  | 1:30000 | Sigma-Aldrich | Immunofluorescence |
| MitoTracker |  | 1µM | Invitrogen | Flow cytometry |
| ROS Brite 670 |  | 5µM | Biomol; AAT Bioquest | Flow cytometry |
| Tetramethylrhodamine (TMRM) |  | 100nM | Thermo Fisher Scientific | Flow cytometry |
| Sarcomeric α-actinin | mouse IgG1 | 1:800 | Sigma-Aldrich | Flow cytometry |
| MYH1E (MF20) | mouse IgG2a | 1:50 | Hybridoma Bank, University of Iowa, US | Flow cytometry |
| Troponin-T | mouse IgG1 | 1:100 | Richard Allan Scientific | Flow cytometry |
| Anti-mouse IgG Alexa Fluor 647 | Donkey | 1:300 | Jackson ImmunoResearch | Flow cytometry |

---

**Data S1:**

**Table S1: Variants detected by mtDNA sequencing.** List of all variants detected by mtDNA sequencing of 26 iPSC clones at early passage (on average p6.5), of additional 7 of the iPSC clones at intermediate (p30) and high (p50) passage, of the 10 corresponding parental cell populations, and cardiomyocytes differentiated off 4 of the iPSC clones during differentiation process at differentiation day (dd) 0, dd5, and dd15. Variants were divided into groups and categories: Variants of group A, non-transmitted variants, were detected only in parental cell populations. Group B variants were homoplasmic in parental cell population and iPSCs derived thereof. Group C, heteroplasmic variants were detected in an iPSC clone with AF > 0.02 at any passage but were less frequent in the corresponding parental cell population. A final group comprises variants found heteroplasmic in differentiated cardiomyocytes. Those variants basically belong to the group of heteroplasmic variants that were transmitted to cardiomyocytes during differentiation. Variants of category 1 are haplotype variants defining a haplogroup. Category 2 variants are polymorphisms and defined by a population variant frequency (MITOMAP (NCBI GenBank)) ≥ 0.02. Category 3 variants are donor-specific variants which were unique to a donor and rare in the population context. heteroplasmic variants were further divided into the groups of amplified variants (which increase in heteroplasmy level of > 10 fold during iPSC culture expansion), stable variants (with fold change ~1), purified variants (decreased by > 10 fold), fluctuating variants (change of heteroplasmy level did not allow confident classification in one of the above mentioned groups), and “nd in p30-50” variants (iPSC clones not analyzed in high passages). Identification of haplotype variants and polymorphisms was based on HaploGrep, NCBI GenBank variant frequency (obtained from MITOMAP), and HmtVar. Variant effect prediction was performed, based on a consensus of in silico prediction algorithms obtained from snpEff impact, CADD, Condel, and HmtVar, and putatively actionable variants are marked in red. A choice of variants with very low variant allele frequencies in iPSC clones and/or parental cells were additionally investigated individually within their genetic context by inspecting and counting sequencing reads directly. Variant allele frequencies that were determined using this method are underlined in yellow. (); existence of variant in parental cell population or iPSC clone not confirmed with statistical confidence (p-value > 0.05).

**Table S2a: Manual determination of very low variant allele frequencies of single nucleotide variants (SNVs)**. The table lists SNVs and their allele frequencies in iPSC clones and parental cell population determined by individual examination of variants within their genetic context. Results are displayed as the number of reads per respective nucleotide embedded in an 11bp sequence. The number for reference allele reads are given in bold, and for variant allele reads in red. Average local error rates calculation and results, statistical test results, as well as local detection limits are presented.

**Table S2b: Manual determination of very low variant allele frequencies of small insertions.** The table lists the small insertions and their allele frequencies in iPSC clones and parental cell population determined by individual examination of variants within their genetic context. Results are displayed as the number of reads for the reference (bold) and variant sequence (red) embedded in an 11bp sequence.

**Data S2: Galaxy workflows.**

