## Supplementary material for "iPSC culture expansion selects against putatively actionable mutations in the mitochondrial genome": Data S2: Galaxy mtDNA analysis workflow.txt

      Galaxy Workflow ' mtDNA analysis workflow 2020'

------------------------------------------------------------------------
Step	Annotation

Step 1: Input dataset
R1.fastq

/select at runtime/
	


Step 2: Input dataset
R2.fastq

/select at runtime/
	


Step 3: Input dataset
Homo_sapiens.GRCh38.dna.chromosome.MT.fa

/select at runtime/

	

Step 4: Input dataset
Homo_sapiens.GRCh38.dna.chromosome.MT.fa.fai

/select at runtime/
	


Step 5: FastQC
Short read data from your current history

Output dataset 'output' from step 1
Contaminant list

/select at runtime/
Submodule and Limit specifing file

/select at runtime/
	


Step 6: FastQC
Short read data from your current history

Output dataset 'output' from step 2
Contaminant list

/select at runtime/
Submodule and Limit specifing file

/select at runtime/
	


Step 7: Trim Galore!
Is this library paired- or single-end?

Paired-end
Reads in FASTQ format

Output dataset 'output' from step 1
Reads in FASTQ format

Output dataset 'output' from step 2
Adapter sequence to be trimmed

Illumina universal
Trims 1 bp off every read from its 3' end.

False
Remove N bp from the 3' end of read 1

10
Remove N bp from the 3' end of read 2

30
Trim Galore! advanced settings

Full parameter list
Trim low-quality ends from reads in addition to adapter removal (Enter
phred quality score threshold)

28
Overlap with adapter sequence required to trim a sequence

1
Maximum allowed error rate

0.1
Discard reads that became shorter than length N

50
Instructs Trim Galore! to remove N bp from the 5' end of read 1

Not available.
Instructs Trim Galore! to remove N bp from the 5' end of read 2 (Only
for paired-end reads)

Not available.
Generate a report file

True
specify if you would like to retain unpaired reads

Do not output unpaired reads
RRBS specific settings

Use defaults (no RRBS)
	


Step 8: FastQC
Short read data from your current history

Output dataset 'trimmed_reads_pair1' from step 7
Contaminant list

/select at runtime/
Submodule and Limit specifing file

/select at runtime/


	
Step 9: FastQC
Short read data from your current history

Output dataset 'trimmed_reads_pair2' from step 7
Contaminant list

/select at runtime/
Submodule and Limit specifing file

/select at runtime/


	
Step 10: Map with BWA-MEM
Will you select a reference genome from your history or use a built-in
index?

Use a genome from history and build index
Use the following dataset as the reference sequence

Output dataset 'output' from step 3
Algorithm for constructing the BWT index

Auto. Let BWA decide the best algorithm to use
Single or Paired-end reads

Paired
Select first set of reads

Output dataset 'trimmed_reads_pair1' from step 7
Select second set of reads

Output dataset 'trimmed_reads_pair2' from step 7
Enter mean, standard deviation, max, and min for insert lengths.

Empty.
Set read groups information?

Do not set
Select analysis mode

1.Simple Illumina mode


	
Step 11: Stats
BAM file

Output dataset 'bam_output' from step 10
Minimum coverage

1
Maximum coverage

10000
Coverage step

10
Exclude reads marked as duplicates

False
Output

One single summary file
Filter by SAM flags

Do not filter
Size of GC-depth bins

20000.0
Maximum insert size

8000
Minimum read length to generate statistics for

Not available.
Report only the main part of inserts

0.99
BWA trim parameter

0
Use reference sequence

Do not use reference


	
Step 12: FastQC
Short read data from your current history

Output dataset 'bam_output' from step 10
Contaminant list

/select at runtime/
Submodule and Limit specifing file

/select at runtime/


	
Step 13: FreeBayes
Choose the source for the reference genome

History
BAM file

Output dataset 'bam_output' from step 10
Use the following dataset as the reference sequence

Output dataset 'output' from step 3
Limit variant calling to a set of regions?

Do not limit
Choose parameter selection level

5. Full list of options
Additional inputs

Do not provide additional inputs
Reporting options

Use defaults
Population model options

Set population model options
The expected mutation rate or pairwise nucleotide diversity among the
population under analysis

0.001
Set ploidy for the analysis

10
Assume that samples result from pooled sequencing

False
Output all alleles which pass input filters, regardles of genotyping
outcome or model

False
Reference allele options

Use defaults
Allelic scope options

Use defaults
Turn off left-alignment of indels

False
Input filters

Set input filters
Include duplicate-marked alignments in the analysis.

False
Exclude alignments from analysis if they have a mapping quality less than

1
Exclude alleles from analysis if their supporting base quality less than

0
Consider any allele in which the sum of qualities of supporting
observations is at least

0
Consider any allele in which and the sum of mapping qualities of
supporting reads is at least

0
Mismatch filters

No mismatch filters (default)
Exclude reads with more than this number of separate gaps

1000
Use stringent input base and mapping quality filters

False
Require at least this fraction of observations supporting an alternate
allele within a single individual in the in order to evaluate the position

0.02
Require at least this count of observations supporting an alternate
allele within a single individual in order to evaluate the position

2
Require at least this sum of quality of observations supporting an
alternate allele within a single individual in order to evaluate the
position

0
Require at least this count of observations supporting an alternate
allele within the total population in order to use the allele in analysis

1
Require at least this coverage to process a site

100
Population and mappability priors

Use defaults
Genotype likelihood options

Use defaults
Algorithmic features

Use defaults
	
